# Druggable CSNK2A1-MeCP2 regulatory axis links Okur-Chung neurodevelopmental syndrome with *MECP2*-related disorders

**DOI:** 10.64898/2026.11.05.764603

**Authors:** Ashley G. Anderson, Srinivas Chamakuri, Rebecca Meyer-Schuman, Elizabeth Hai-Yen Chu, Jean-Pierre Revelli, Jiyoen Kim, Manar Zaghlula, Surabi Veeraragavan, Damian W. Young, Huda Y. Zoghbi

**Affiliations:** Department of Molecular and Human Genetics, Baylor College of Medicine, Houston, TX; Jan and Dan Duncan Neurological Research Institute at Texas Children’s Hospital, Houston, TX; Center for Drug Discovery, Baylor College of Medicine, Houston, TX 77030; Department of Pathology & Immunology, Baylor College of Medicine, Houston, TX 77030; Innovative Genomics Institute, University of California, Berkeley, Berkeley, CA; Verna and Marrs McLean Department of Biochemistry and Molecular Pharmacology, Baylor College of Medicine, Houston, TX; Department of Pediatrics, Baylor College of Medicine, Houston, TX; Howard Hughes Medical Institute, Baylor College of Medicine, Houston, TX; Department of Neuroscience, Baylor College of Medicine, Houston, TX

## Abstract

The brain is exquisitely sensitive to methyl-CpG-binding protein 2 (MeCP2) abundance. Too little causes Rett syndrome (RTT), too much causes *MECP2* duplication syndrome (MDS), and in both, severity tracks with the amount of functional MeCP2. Even modest deviations are detrimental, yet how MeCP2 abundance is controlled remains poorly understood. Using genome-wide CRISPR knockout screens, we identified hundreds of proteins regulating MeCP2, revealing a casein kinase 2 alpha 1 (CSNK2A1)-driven regulatory axis stabilizing MeCP2. We show this axis is druggable and therapeutically relevant, using a novel CSNK2A1 inhibitor in human cells and targeting *Csnk2a1* in an MDS mouse model. We further connect CSNK2A1-MeCP2 regulatory disruption to Okur-Chung neurodevelopmental syndrome (OCNDS), caused by mutations in *CSNK2A1*, finding ∼50% less hippocampal MeCP2 in an OCNDS mouse model. Together, these data identify a druggable regulatory axis modulating MeCP2 and implicate MeCP2 dysregulation in the molecular pathogenesis of OCNDS and potentially other neurodevelopmental disorders.

---

Dosage-sensitive genes have been implicated in a number of neurodevelopmental (NDD) and neuropsychiatric disorders, including autism spectrum disorder (ASD), attention-deficit/hyperactivity disorder (ADHD), and schizophrenia^1–3^. Many of these disease-linked, dosage-sensitive genes encode proteins involved in chromatin binding and transcriptional regulation pathways, exemplified by the X-linked gene methyl-CpG-binding protein 2 (*MECP2*)^3–6^. Loss-of-function mutations in *MECP2* cause Rett syndrome (RTT), a severe neurological disorder characterized by a period of apparently normal early development followed by regression of motor and language skills, stereotypic hand movements, ataxia, intellectual disability, seizures, and progressive motor decline^7,8^. Conversely, increased *MECP2* dosage caused by duplications or triplications of the *MECP2* locus results in *MECP2* duplication syndrome (MDS), an equally devastating neurological disorder that shares clinical similarities to RTT but differs in sex bias and disease progression^9–11^. Whereas RTT typically affects girls and arises via *de novo MECP2* mutations, MDS primarily affects boys and is often inherited from carrier mothers^10^. Genotype-phenotype studies, together with profiling of disease mouse models, show a correlation between the phenotypic severity of both disorders and the amount of MeCP2 expression or function remaining in the cell, with complete deletion (no MeCP2) or triplications of *MECP2* (3X more MeCP2) being the most severe^12–17^. Even a modest ∼30% deviation from normal MeCP2 levels is sufficient to cause neuropsychiatric phenotypes^18^. Despite the importance of precisely regulating MeCP2 for normal brain function, how MeCP2 protein abundance is set and maintained within such as narrow range remains poorly understood.

Modifying levels of MeCP2 has proven to be therapeutic in both mouse models and patient derived iNeurons of RTT and MDS, strongly supporting the design of therapeutic strategies to directly modulate MeCP2 abundance^19–23^. Clinical trials for *MECP2*-related disorders are actively pursuing this strategy, including AAV-based gene therapies to restore functional *MECP2* in RTT and ASO-treatments to reduce *MECP2* in MDS. Although these approaches are promising, identifying alternative ways to modulate MeCP2 stability could expand non-invasive therapeutic options^24^ and complement existing strategies to enable precise modulation of MeCP2 levels.

MeCP2 is highly expressed in neurons^25–27^ and binds methylated cytosines broadly^28–30^, particularly during postnatal development, where it interacts with large protein complexes to regulate gene expression^31,32^. Notably, the molecular signatures of RTT and MDS are mirror images of each other– i.e. genes that are upregulated in RTT are downregulated in MDS^33^. These transcriptional changes are also highly sensitive to changes in the abundance of MeCP2 ^20,34^. Although many downstream consequences of MeCP2 function and critical domains of the protein have been well-defined, our understanding of the upstream mechanisms that regulate MeCP2 remains incomplete. Understanding these regulatory pathways could advance therapeutic development for *MECP2*-related disorders and reveal molecular pathways potentially relevant to other NDDs.

To uncover pathways regulating MeCP2, we performed a series of CRISPR knockout screens to identify genes involved in stabilizing or destabilizing MeCP2 and identified ∼1800 putative modulators of MeCP2 levels. Many MeCP2 regulators were themselves NDD risk genes, including casein kinase 2 alpha 1 (CSNK2A1), a screen hit stabilizing MeCP2 that we prioritize for further study. CSNK2A1 is a ubiquitous, brain-enriched serine/threonine kinase that encodes the catalytically active subunit of the well-characterized CK2 holoenzyme, composed of both alpha (CSNK2A1 or CSNK2A2) and beta (CSNK2B) subunits^35,36^. Heterozygous, *de novo* mutations in *CSNK2A1* cause Okur-Chung Neurodevelopmental Syndrome (OCNDS)^37,38^, a severe neurological disorder that shares clinical features with *MECP2*-related disorders, including neurodevelopmental delay, autism, loss of language, sleep disruption, motor coordination deficits, microcephaly, seizures, and stereotypies^10,39^. *CSNK2B* mutations are also associated with a severe epileptic NDD, known as Poirier-Bienvenu neurodevelopmental syndrome^40^. Based on our screen results and this human genetic evidence, we focused our downstream analyses on understanding the regulatory relationship between CSNK2A1 and MeCP2.

Here, we show that CSNK2A1 interacts with MeCP2 throughout mouse brain development and that inhibiting CSNK2A1 kinase activity destabilizes MeCP2 via a neddylation-dependent ubiquitin proteasomal pathway. We further demonstrate that genetic reduction of *Csnk2a1* in the *MECP2^TG1^* mouse model of MDS moderately lowers MeCP2 levels and selectively rescues locomotor activity and repetitive rearing behavior, while partially rescuing the MDS transcriptional disease signature, including core MeCP2-regulated genes. Finally, we shed light on the molecular pathogenesis of OCNDS by finding significantly lower MeCP2 levels specifically in the hippocampus of *Csnk2a1*^K198R/+^ mice, a mouse model harboring the most common mutation found in the disorder. Together, these data advance our molecular understanding of how MeCP2 protein dosage is regulated and identify druggable pathways for modulating MeCP2 with therapeutic implications for *MECP2*-related disorders, OCNDS, and potentially other NDDs.

## RESULTS

### CRISPR KO screens reveal post-translational regulators of MeCP2

To examine post-translational regulation of MeCP2, we used the global protein stability reporter system (GPS) in combination with two CRISPR KO screens to identify protein-coding genes important for either stabilizing or de-stabilizing MeCP2 ^41^. The GPS system allows for the expression of an internal control (dsRed) and MeCP2 fused to an enhanced green fluorescent protein (eGFP) from the same bicistronic mRNA transcript. We generated a clonal human embryonic kidney (HEK)-293T cell line that stably expressed Cas9 and the bicistronic vector *dsRED-IRES-MECP2-EGFP* (**Figure 1A**). We validated this cell line using several methods to confirm Cas9 functionality and to measure the efficiency of targeting MeCP2 directly using single guide RNA (sgRNAs) against either *eGFP* or *MECP2* (**Supplemental Figure 1A**). Using this cell line, we could use fluorescently activated cell sorting (FACS) to separate cells using their GFP/dsRed ratio into low MeCP2 expressing cells (bottom 10% of GFP/dsRed ratio, low group) or high MeCP2 expressing cells (top 10% of GFP/dsRed ratio, high group), followed by sgRNA amplification, NGS sequencing, and sgRNA enrichment analysis (**Figure 1A**). We performed sequential CRISPR-Cas9 enrichment screens using distinct sgRNA libraries targeting: (1) a smaller library of druggable kinase-associated genes (kinome, 763 kinase-domain containing genes) and (2) most protein-coding genes in human genome (genome-wide, 19,114 genes) (**Figure 1A**). While the genome-wide library also targets genes contained in the kinome library, we started our screen with the kinome library for the following reasons: 1) the kinome is a smaller library with higher power (8 sgRNAs/gene) to detect significant, small changes to MeCP2 levels, compared to the genome-wide screen (4 sgRNAs/gene), 2) we reasoned that prioritizing overlapping hits from both screens would identify robust, druggable MeCP2 regulators.

**Figure 1.**
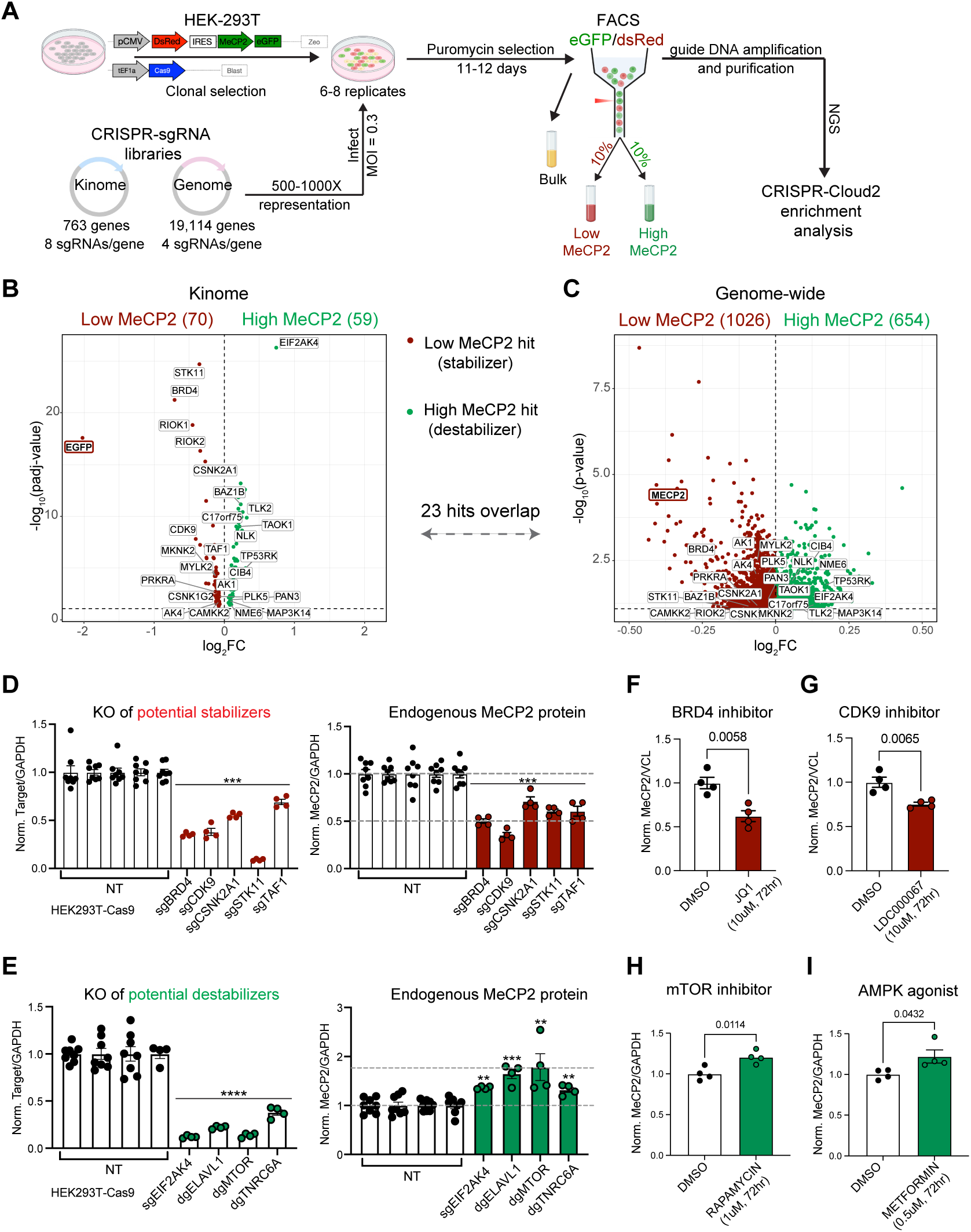
CRISPR KO screens identify new regulators of MeCP2 stability. A) Schematic of the CRISPR screen workflow. B-C) Volcano plots of (B) the kinome screen results showing the log_2_FC vs −log_10_ p-adjusted value (FDR<0.05) enrichment score per gene or (C) the genome-wide screen results showing the log_2_FC vs −log_10_ p-value (p<0.08) enrichment score per gene that increased MeCP2 (green) or decreased MeCP2 (red) expression. D-E) Validation of top screen hits on endogenous MeCP2 levels relative to loading control (GAPDH) in HEK293T cells using independent sgRNAs or dual guide RNA vectors (dgRNAs). Combined bar plot showing target gene knockout (KO) efficiency (left) and corresponding changes to MeCP2 levels (right) as measured by western blot quantification compared to respective non-targeting (NT) control samples, stabilizers (red) and destabilizers (green). Data presented as mean ± SEM, **p<0.005, ***p<0.0005, ****p<0.0001 using a one-way ANOVA with post-hoc Dunnett’s multiple comparisons between gene-targeting sgRNAs or dgRNAs to respective NT samples. F-I) Quantification of endogenous MeCP2 levels relative to loading controls (GAPDH or VCL) in HEK293T cells after 72hr incubation with (F) BRD4 inhibitor JQ1 at 10uM, (G) CDK9 inhibitor LDC000067 at 10uM, (H) MTOR inhibitor Rapamycin at 1 uM, or (I) agonist of AMPK pathway Metformin at 0.5uM. Data presented as mean ± SEM, p-values determined using a Student’s t-test.

Following CRISPR Cloud2 enrichment analyses^42^, we identified 129 significant hits from the kinome screen (**Figure 1B**) and 1680 significant hits from the genome-wide screen (**Figure 1C**) that were enriched in either the low-MeCP2 (stabilizers) or high-MeCP2 groups (destabilizers) relative to an unsorted bulk population. 23 kinase-associated hits overlapped in both screen, 21 hits were in the same groups across both screens (**Supplemental Table 1**). Each library contained internal positive controls targeting MeCP2 directly that were significantly enriched in the low group as expected (guides against *GFP* in kinome library or against *MECP2* in the genome-wide library) (**Figure 1B-C**). Several top hits overlapped with validated genes from a previous siRNA arrayed screen^24^, including *PPP2R1A* (genome), *RIOK1* (kinome), *RIOK2* (genome/kinome), *CDK7* (kinome), *JAK1* (kinome), and *ACVR2B* (kinome). *PPP2R1A* knockdown (KD) was shown to decrease MeCP2 *in vivo* and inhibitors of PPP2R1A in a mouse model of MDS lowered levels and rescued motor-learning skills^24^. We also found *CAMK2A* and *CAMK2B*, kinases known to phosphorylate MeCP2 during neural activity, as stabilizers in our whole genome screen. STRING analyses of the top 100 most enriched up or down genome-wide screen hits had 2-4.5fold more protein-protein interactions (PPIs) than expected by chance (p<1.0e^-16^) indicating screen hits are highly interconnected.

Given the positive controls and strong pathway convergence, we next proceeded to validate new candidate regulators of MeCP2 by prioritizing hits that overlapped in both screens and testing whether KO of these targets using independent sgRNAs altered endogenous MeCP2 levels (**Figure 1D-E**). KO of *BRD4*, *CDK9*, *CSNK2A1*, *STK11*, and *TAF1* significantly lowered target protein levels and decreased endogenous MeCP2 (**Figure 1D**), whereas KO of *EIF2AK4* significantly lowered target protein levels and increased MeCP2 levels (**Figure 1E**), consistent with the directionality of the primary screen. We also validated whole-genome hits that altered endogenous MeCP2, including *MTOR, ELAVL1* and *TNRC6A*, genes associated with RNA-binding and splicing, respectively (**Figure1E**). Notably, several validated hits were directionally concordant within known regulatory modules, including promoter-proximal transcription (*BRD4*, *CDK9*, and *TAF1*)^43^ that decreased MeCP2 and nutrient- and energy-sensing signaling (*MTOR* and *STK11*)^44^, where loss of MTOR increased MeCP2 while loss of STK11, an upstream AMPK activator, decreased levels.

To broadly examine whether any hits from the whole-genome screen were enriched in distinct molecular pathways, we performed gene ontology (GO) analyses and found hits that stabilize MeCP2 were enriched in categories of RNA processing and ribosome biogenesis, and hits that destabilize MeCP2 were enriched in categories associated with the mitochondria and electron transport chain (**Supplemental Figure 1B-C**). We also examined brain region and cell-type expression of screen hits using the comprehensive single-cell-RNA-seq dataset from the Allen brain atlas^45^ and found most hits were highly expressed in neural tissue, particularly the overlap of genome-kinome hits, including validated *BRD4*, *CSNK2A1*, *STK11*, and *EIF2AK4* (**Supplemental Figure 1D-E**). As several validated hits are themselves associated with autism spectrum disorder (ASD) and NDDs (*BRD4*, *CSNK2A1*, *TAF1*, and *MTOR*), we asked whether our genome-wide screen hits were enriched for ASD and NDD-associated genes. Using a recently published inheritance-specific (mono-vs bi-allelic) machine learning NDD prediction model that predicts genes implicated in autism spectrum disorder (ASD), developmental delay (DD), or developmental and epileptic encephalopathy (DEE)^46^, we found only modest enrichment for predicted NDD risk genes, with only bi-allelic DD risk genes reaching significance amongst MeCP2 destabilizers (**Supplemental Figure 1F**).

To identify potential drug-gene interactions, we used the Drug Gene Interaction Database (DGIdb) to examine the percent of known U.S Food and Drug Administration (FDA) approved (Tier 1), investigational (Tier 2), or no information (Tier 3) drugs that target proteins from our screen hits. We found ∼16-18% of genome-wide hits and ∼40% of kinase-hits had FDA approved drug targets, revealing a proportion of potentially druggable MeCP2 regulators for future studies (**Supplemental Figure 1G, Supplemental Table 2**). We further tested whether available drugs that either inhibited or activated validated screen hits could also modulate endogenous MeCP2 levels. We found BRD4 (JQ1) and CDK9 (LDC000067), inhibitors significantly lowered MeCP2 levels in HEK293T cells, similar to KO results (**Figure 1F-G**). MTOR inhibitor (Rapamycin) significantly increased endogenous MeCP2 levels, similar to KO results (**Figure 1H**). An activator of the AMPK signaling pathway (Metformin), downstream of STK11, increased endogenous MeCP2 levels, opposite of *STK11* KO, indicating bi-directional modulation of MeCP2 levels targeting the STK11-signaling cascade (**Figure 1I**). Together, these data reveal druggable molecular pathways regulating MeCP2 for further study.

### CSNK2A1 binds to MeCP2 across mouse brain development and stabilizes MeCP2 *in vivo*

To further prioritize top screen hits for therapeutic relevance, we focused on the overlapping kinase-associated genes in both screens that were highly expressed in the brain and asked whether any hits directly interacted with MeCP2 in neuronal tissue, allowing us to prioritize hits with fewer intermediaries and minimize off-target modulatory effects (**Figure 2A**). We overlapped screen hits with published MeCP2 PPI datasets from MeCP2 proximity-labeling in hippocampal neurons^47^ and co-immunoprecipitation (co-IP) followed by mass spectrometry (MS) in cortical neurons^48^ and found two MeCP2-stabilizing hits that overlapped across all datasets, *CSNK2A1* and *PRKRA* (**Figure 2A**, **Supplemental Table 1**). Notably, both genes are linked to human disease. Homozygous mutations in *PRKRA* lead to dystonia 16 (DYT16)^49^, which is characterized by early-onset generalized dystonia. Heterozygous mutations in *CSNK2A1*, encoding the catalytic alpha subunit of the CK2 kinase, cause Okur Chung Neurodevelopmental syndrome (OCNDS), characterized by developmental delay, intellectual disability, loss of language, microcephaly, ASD/ADHD, and motor coordination impairments, and stereotypies^37^. We were particularly interested in *CSNK2A1* given the overlapping features of OCNDS and *MECP2*-related disorders (**Figure 2B**), high brain expression, and druggability.

**Figure 2.**
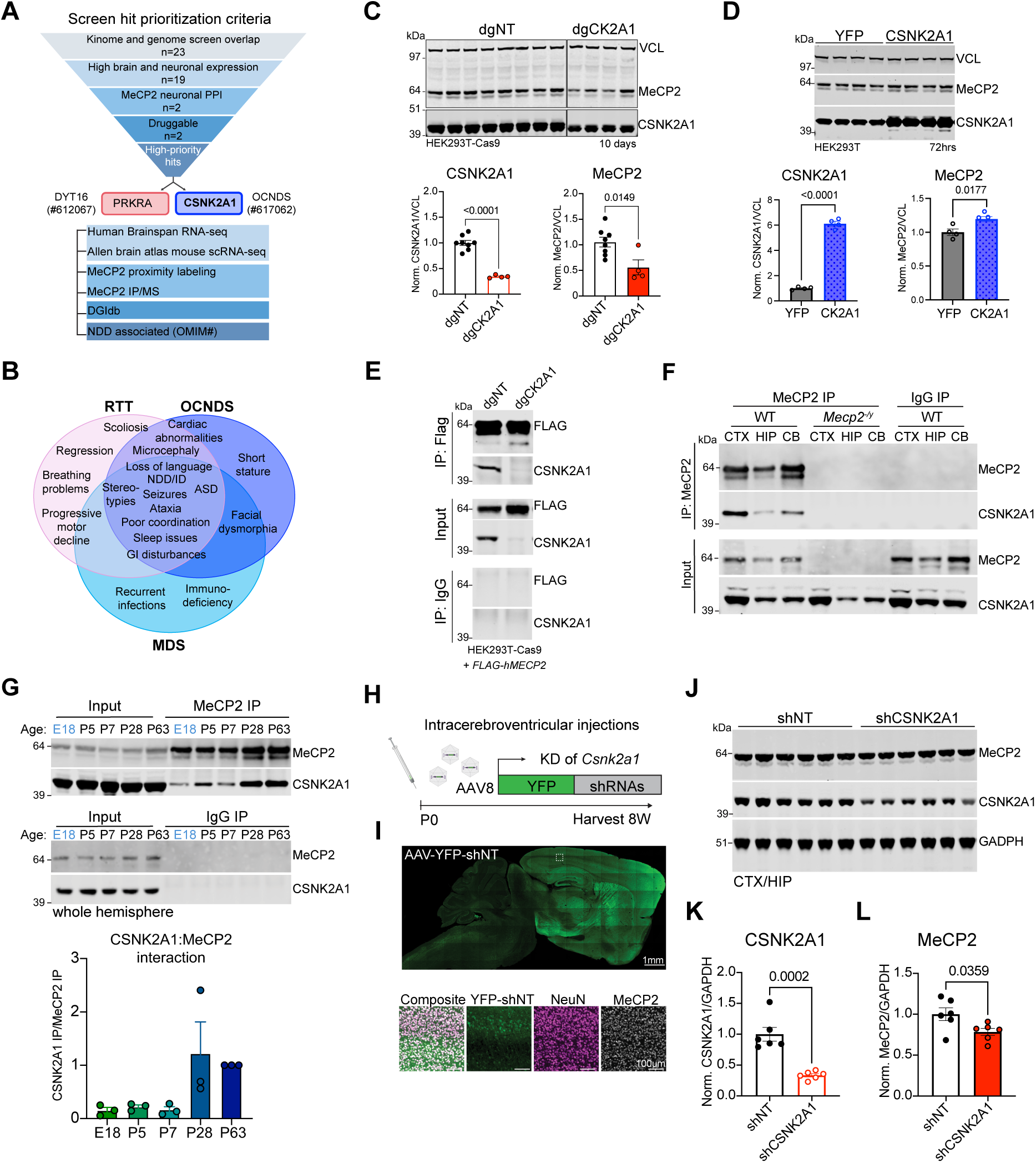
CSNK2A1 interacts with MeCP2 and modulates its stability *in vivo*. A) Prioritization of screen hits filtered by overlapping kinome-genome, human and mouse brain expression, MeCP2 PPIs^47,48^, druggability, and disease association (OMIM#). B) Venn diagram of overlapping clinical phenotypes of Rett syndrome (RTT), Okur Chung Neurodevelopmental Syndrome (OCNDS), and *MECP2* duplication syndrome (MDS). C) Western blot and quantification of CSNK2A1 and MeCP2 total protein levels relative to loading control (VCL) in HEK293T-Cas9 cells infected with either dgCK2A1 or dgNT KO. D) Western blot and quantification of CSNK2A1 and MeCP2 levels relative to loading control (VCL) from HEK-293Ts overexpressing either YFP or CSNK2A1. Data presented as mean ± SEM, p-values calculated using a student’s t-test for C-D. E) IP western blot for FLAG or IgG in CSNK2A1 KO or WT (NT) HEK293Ts overexpressing Flag-MeCP2, probing for FLAG and CSNK2A1 in input and IP samples. F) IP western blot of endogenous MeCP2 or IgG control from mouse brain regions: cortex (CTX), hippocampus (HIP), or cerebellum (CB) in WT or *Mecp2^-/y^* (MeCP2 null) mice. G) IP of MeCP2 at embryonic (E18) and post-natal timepoints (P5, P7, P28, and P63) and quantification of MeCP2-CSNK2A1 interaction over development (representative western blot from replicate 1). H) Schematic of intracerebroventricular (ICV) injection protocol using an AAV8 miR30-YFP 4-shRNA virus targeting *Csnk2a1* or NT into P0 WT pups. I) 20X immunofluorescent (IF) image of the viral spread post-injection of an AAV-YFP-shNT injection at 4 weeks and colocalization with MeCP2 (gray) and neuron marker NeuN (magenta). J) Western blot and quantification of combined cortical/hippocampal protein samples at 8 weeks post-injection measuring levels of (K) CSNK2A1 and (L) MeCP2 relative to loading control (GAPDH). Data presented as mean ± SEM, p-values calculated using a student’s t-test between shNT and shCSNK2A1 samples.

To further validate *CSNK2A1*, we used a dual sgRNA vector to lower CSNK2A1 levels by over 70% at both the protein and RNA levels (**Figure 2C, Supplemental Figure 2A**) and confirmed a decrease of endogenous MeCP2 protein with no change to *MECP2* transcript levels (**Figure 2C**, **Supplemental Figure 2B**). Conversely, overexpression of CSNK2A1 in human cells significantly increased MeCP2 levels relative to YFP control, showing that MeCP2 can be bi-directionally modified by CSNK2A1 activity (**Figure 2D**). To confirm the interaction between MeCP2 and CSNK2A1, we performed a co-IP experiment by pulling down Flag-tagged MeCP2 overexpressed in either WT (dgNT) or CSNK2A1 KO (dgCSNK2A1) HEK293T cells and found that CSNK2A1 interacted with MeCP2 in WT cells, but not in CSNK2A1 KO cells (**Figure 2E**).

We next asked if this interaction was preserved *in vivo* and performed co-IP experiments for endogenous MeCP2 from either WT or *Mecp2^-/y^* (*Mecp2* null) mouse brain regions (cortex, hippocampus, and cerebellum). Again, we found that MeCP2 interacted with CSNK2A1 across all brain regions tested in WT samples, but not in *Mecp2* null or control IgG samples from the same WT tissue (**Figure 2F**). This interaction was preserved across mouse brain development from late embryonic (E18) and early postnatal timepoints to adulthood (P63), peaking at 4 weeks (P28), a timepoint that coincides with postnatal stabilization of MeCP2 levels (**Figure 2G, Supplemental Figure 2C-D**). We also examined the co-expression pattern of *Csnk2a1* and *Mecp2* throughout the mouse brain using dual fluorescent *in situ* hybridization and found they were ubiquitously co-expressed across all brain regions (**Supplemental Figure 2E-G**). A notable cell-type specific difference in expression pattern was observed in the cerebellum, where *Csnk2a1* had greater expression in the Purkinje layer compared to *Mecp2* expression that was distributed evenly across Purkinje and granular layers (**Supplemental Figure 2F**), which was also observed in adult mouse single-cell RNA-seq data (**Supplemental Figure 1E**).

Since both MeCP2 and CSNK2A1 are ubiquitously co-expressed, we next tested whether knockdown (KD) of *Csnk2a1* could alter MeCP2 stability *in vivo*. We injected an adeno-associated virus (AAV) short hairpin RNA (shRNA) virus fused to yellow fluorescent protein (YFP) targeting *Csnk2a1* or non-targeting into the ventricles of wild-type, post-natal day 0 (P0) mouse pups (**Figure 2H**). Wide viral spread in neuronal cell-types was detected using this method throughout the adult brain by imaging the YFP signal and co-localization with neuronal marker NeuN and MeCP2 (**Figure 2I**). KD of *Csnk2a1* (**Figure 2J-K, Supplemental Figure 2H)** significantly lowered MeCP2 protein but not *Mecp2* RNA levels (**Figure 2L, Supplemental Figure 2I**). These data show that CSNK2A1 modulates MeCP2 stability *in vivo* and could be a tractable target to lower MeCP2 in MDS.

### Novel inhibitor of CSNK2A1 reduces MeCP2 stability via neddylation-dependent ubiquitin-proteasomal pathway

We next asked whether inhibiting CSNK2A1 activity could alter MeCP2 protein levels and tested the known CK2 inhibitor, TBB, in HEK293T cells. We found only a minimal ∼15% change in MeCP2 at the highest dose (**Figure 3A).** Testing another CSNK2A1 inhibitor, CX-4945 (Silmitasertib), at the same conditions showed signs of cellular toxicity, but lower doses of CX-4945 (250nM) decreased MeCP2 levels by ∼25% (**Supplemental Figure 3A**). Given these promising results, we next examined a novel CSNK2A1 inhibitor, CDD-2021 (Ki=14nM for CSNK2A1, 62nM for CSNK2A2) identified from DNA-encoded chemical libraries as previous described^50–53^(**Figure 3B**, **Supplemental chemistry methods**). We found that CDD-2021 lowered MeCP2 levels by ∼30% with no obvious cellular toxicity (**Figure 3C**). This decrease was dose dependent at low concentrations with no effect on *MECP2* RNA levels, demonstrating that its biological effect is specific and controllable rather than random or off-target (**Figure 3C**). We further tested the effect of CSNK2A1 inhibition on MeCP2 protein stability using a doxycycline-inducible (DOX) system overexpressing WT-MeCP2 +/− TBB, CDD-2021, or DMSO (vehicle) (**Figure 3D**). We found that CDD-2021 significantly reduced MeCP2 stability over the course of three day compared to DMSO, whereas no effect was observed with TBB (**Figure 3D**).

**Figure 3.**
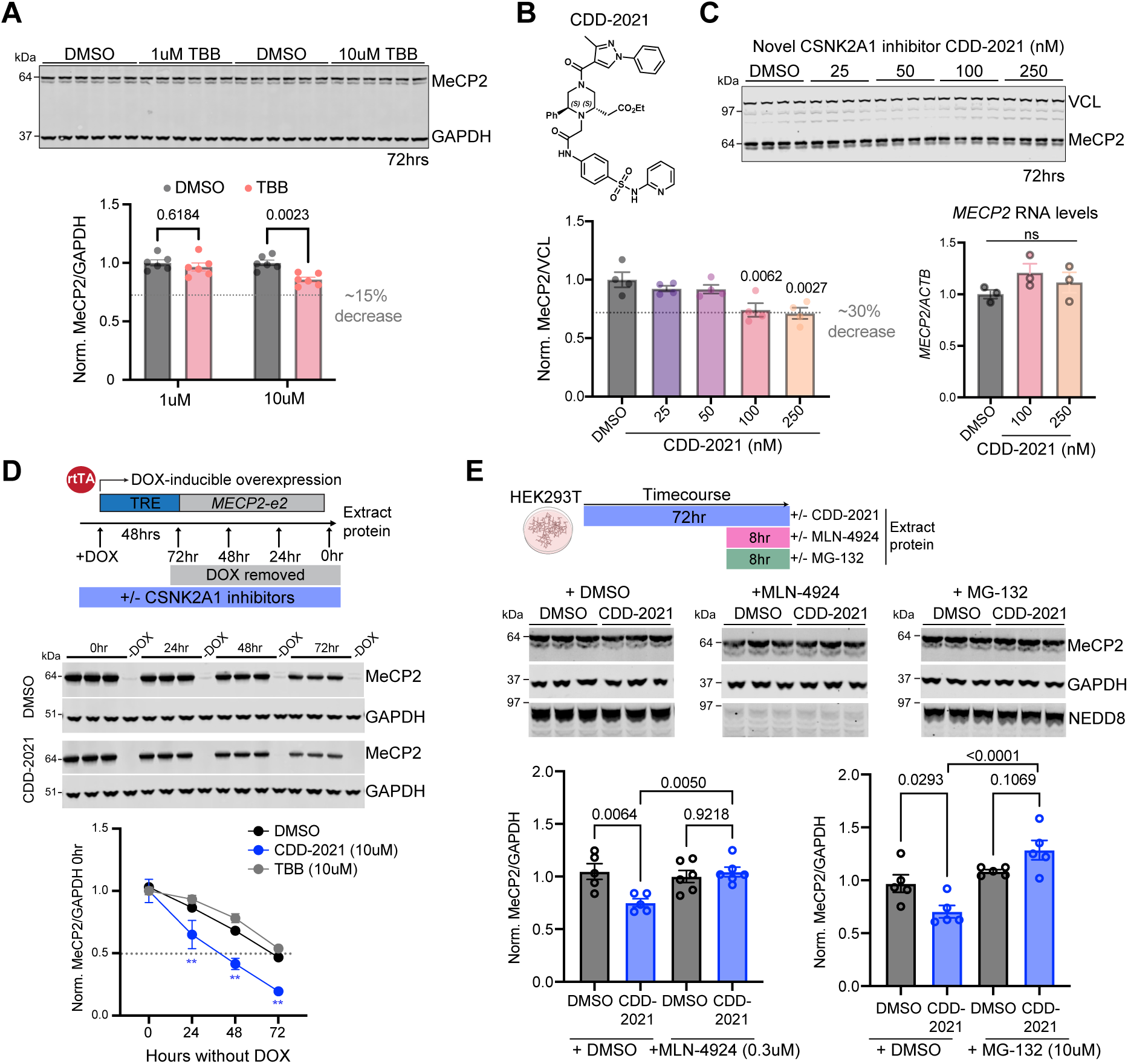
Novel CSNK2A1 inhibitor reduces MeCP2 stability via a neddylation-dependent ubiquitin-proteasomal pathway. A) Western blot and quantification of MeCP2 protein levels relative to loading control (GAPDH) in HEK293T cells incubated with CSNK2A1 inhibitor TBB or DMSO vehicle control for 72hrs at 1uM or 10uM. B) Structure of CDD-2021 compound and (C) western blot and quantification of MeCP2 protein levels (left) relative to loading control (VCL) or *MECP2* RNA levels relative to *ACTIN* (housekeeping gene) via qRT-PCR (right) in HEK293T cells incubated with CDD- 2021 at nM doses for 72hrs. Data presented as mean ± SEM, p-values determined using a one-way ANOVA with post-hoc Dunnett’s multiple comparisons. D) Schematic of experimental design. Western blots and quantification of MeCP2 stability relative to loading control (GAPDH) using the DOX-inducible experimental design to overexpress MeCP2-e2 in the presence of TBB (10uM), CDD2021 (10uM), or DMSO (control). Data presented as mean ± SEM, p-values determined using a two-way ANOVA with Tukey’s post-hoc multiple comparisons **P<0.005. E) Timeline of experimental design. Western blots and quantification of endogenous levels of MeCP2 relative to loading control (GAPDH) following treatment with DMSO or CDD-2021 (250nM) for 72hrs followed by incubation with either MLN-4924 (0.3uM, Nedd8-activating enzyme inhibitor), MG-132 (10uM, proteasome inhibitor), or DMSO (vehicle) 8hrs prior to harvest. NEDD8 blot shows MLN-4924 effectively inhibits neddylation. Data presented as mean ± SEM, p-values determined using a one-way ANOVA with post-hoc Sidak’s multiple comparisons.

We next asked how CSNK2A1 inhibition via CDD-2021 causes MeCP2 degradation. Previous studies have implicated the ubiquitin-proteasomal system (UPS) in MeCP2 degradation^21,54^ and several ubiquitin-proteasomal associated genes appeared on our genome-wide screen, including adapters for the Cullin-RING finger E3 ligases (CRLs). Members of the CRL family, such as CUL3, are also associated with NDDs^55^. To test whether CRLs mediate the CDD-2021 degradation of MeCP2, we tested whether inhibiting the CRL-dependent neddylation degradation pathway via the neddylation activating enzyme inhibitor (MLN-4924) could reverse the degradation of MeCP2 in the presence of CDD-2021 (**Figure 3E**). We confirmed that MLN-4924 blocked neddylation via loss of ∼97 kDa neddylated cullin signal and found that MeCP2 levels were restored back to control levels in the presence of CDD-2021 (**Figure 3E**). We also found inhibiting the proteasome (MG-132) fully restored MeCP2 levels (**Figure 3E**). To examine other degradation mechanisms, we also examined the autophagy-lysosome pathway by overexpressing both HA-tagged Ubiquitin (Ub) and FLAG-tagged MeCP2 in HEK293T cells +/−CDD-2021, +/−inhibitors of the lysosome (CQ) or the UPS (MG-132). We found that CDD-2021 reduced MeCP2 ubiquitination and only MG-132, not CQ, blocked this reduction (**Supplemental Figure 3B-D**). This indicates that CDD-2021 increases MeCP2 susceptibility to Ub-mediated degradation via the UPS, not an autophagy-lysosomal pathway. Together, these data show that CSNK2A1 protects MeCP2 from degradation via a CRL-dependent, ubiquitin-proteasomal pathway.

### CSNK2A1 alters phosphorylation levels of distinct MeCP2 serine residues that do not individually influence MeCP2 stability

To examine whether CSNK2A1 can directly phosphorylate MeCP2, we used a recombinant protein system with the ADP-Glo kinase assay. While CSNK2A1 increased the phosphorylation signal of MeCP2 *in vitro*, the signal was mild compared to known positive control casein and CDD-2021 significantly decreased signal for both proteins (**Supplemental Figure 4A**). However, mass spectrometry analysis using recombinant MeCP2 and CSNK2A1 +/−CDD-2021 could not detect any phosphorylation marks on MeCP2 despite 65% coverage (**Supplemental Figure 4B**). These findings suggest that CSNK2A1 alone does not efficiently phosphorylate MeCP2 and that other cellular co-factors might be needed, such as the complete CK2 holoenzyme. We therefore used a cell-based system under physiologically relevant conditions to identify any CSNK2A1-dependent phosphorylation sites on MeCP2. We knocked out *CSNK2A1* using sgRNAs while overexpressing a FLAG-MECP2 construct, followed by immunoprecipitation-mass spectrometry (IP-MS) (**Figure 4A**). We achieved ∼65% coverage of MeCP2 and found several phosphorylated serine residues (S80, S229, S274, S326, and S360), including known activity-dependent phospho-sites sites S80 and S274 (**Figure 4A**). Three serine residues in particular had reduced phosphorylation with KO of *CSNK2A1* at S80, S229, and S326 compared to the non-targeting control (**Figure 4B**). These phosphorylation sites were not specific to a functional protein domain in MeCP2 but spread across the methyl CpG binding domain (MBD), the transcriptional repressor domain (TRD), and the C-terminal domain (CTD) of MeCP2 (**Figure 4A**).

**Figure 4.**
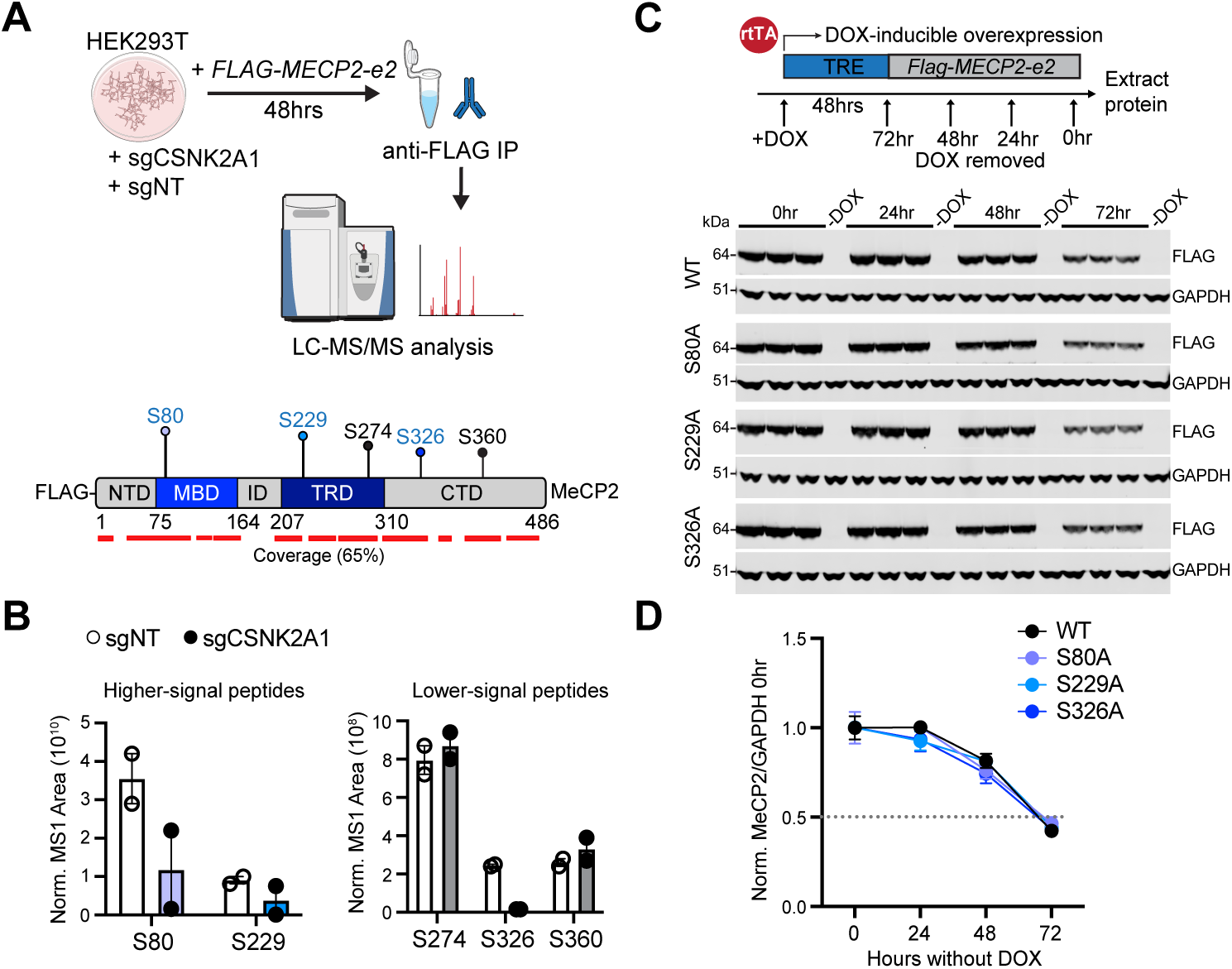
CSNK2A1 alters MeCP2 phosphorylation at distinct serine residues that do not independently alter MeCP2 stability. A) Schematic of the IP/MS experiment and coverage (red line) with location of phosphorylated serine residues in distinct protein domains of MeCP2 (NTD= N-terminal domain, MBD= methyl CpG binding domain, ID= intervening domain, TRD= transcriptional repressor domain, CTD= C-terminal domain). B) Graph showing normalized quantification of phosphorylated peptides over total peptides detected in sgNT and sgCSNK2A1 samples (N=2/condition, no statistical comparisons). C-D) Schematic of pInducer DOX-inducible experiment with *Flag-MECP2* (WT) vs *Flag-MECP2* serine to alanine mutations (S80A, S229A, or S326A) with western blots and quantification of MeCP2 relative to loading control (GAPDH) across each timepoint normalized to 0hr (continuous DOX) timepoint. Data presented as mean ± SEM, p-values determined using a two-way ANOVA with Tukey’s post-hoc multiple comparisons.

To test whether these residues alter MeCP2 stability, we made serine to alanine (S to A) mutations at each site using a doxycycline-inducible (DOX) system^56^ to overexpress FLAG-tagged MECP2 constructs: WT-MECP2, S80A, S229A, or S326A (**Figure 4C**). We then measured the stability of MeCP2 after DOX removal every 24hrs over the course of three days and found none of the single mutations significantly impacted MeCP2 protein stability (**Figure 4D**). These data suggest that: 1) CSNK2A1 likely requires other cellular components to influence MeCP2 phosphorylation and 2) either a combination of phosphorylation sites might be necessary to alter MeCP2 protein stability or CSNK2A1 activity might influence MeCP2 protein stability indirectly via other phospho-dependent mechanisms.

### Genetic reduction of *Csnk2a1* lowers MeCP2 levels and rescues locomotor activity in MDS model

Since we know that even subtle changes to MeCP2 levels can be therapeutic, we next asked whether genetically reducing *Csnk2a1* in a *MECP2* duplication syndrome mouse model (*TG1* mice)^14^ could lower MeCP2 levels and rescue any behavioral phenotypes. *MECP2^TG^*^1^ mice have a transgenic allele driving the expression of a human *MECP2*^14^. To generate *Csnk2a1* heterozygous mice (*Ck2a1^+/−^*), we crossed conditional *Csnk2a1^flox/flox^* mice^57^ to *Sox2-Cre^+/−^* ;*Csnk2a1*^+/+^ female mice ^58^ that have active Cre-recombinase in germline cells to recombine floxed alleles *in utero* independent of inheriting the *Cre* allele. We crossed *Ck2a1^+/−^* and TG1 mice to generate four genotypes for molecular and behavioral characterization: WT, *Ck2a1^+/−^*, *TG1*, and *Ck2a1^+/−^; TG1* (rescue) mice. We then examined expression of CSNK2A1 and MeCP2 across all genotypes, male and female, within distinct brain regions. We confirmed the 50% reduction of CSNK2A1 protein in *Ck2a1^+/−^* and *Ck2a1^+/−^; TG1* brain tissue and the 2-3X overexpression of MeCP2 in *TG1* and *Ck2a1^+/^; TG1* (**Figure 5A-B**). We found that reduction of *Csnk2a1* significantly lowered MeCP2 levels in *Ck2a1^+/−^* and *Ck2a1^+/−^; TG1* brain regions, which was driven by cerebellum and hippocampal samples when examined separately (**Figure 5C**, **Supplemental Figure S5A-D**). This region-specific driven decrease of MeCP2 suggests region-specific regulation as it was not due to residual CSNK2A1 remaining in distinct regions relative to others (i.e. upregulation of the WT allele), since all regions had a comparable 50% reduction of CSNK2A1 in both *Ck2a1^+/−^* and *Ck2a1^+/−^; TG1* samples (**Figure 5B, Supplemental Figure 5E-H**).

**Figure 5.**
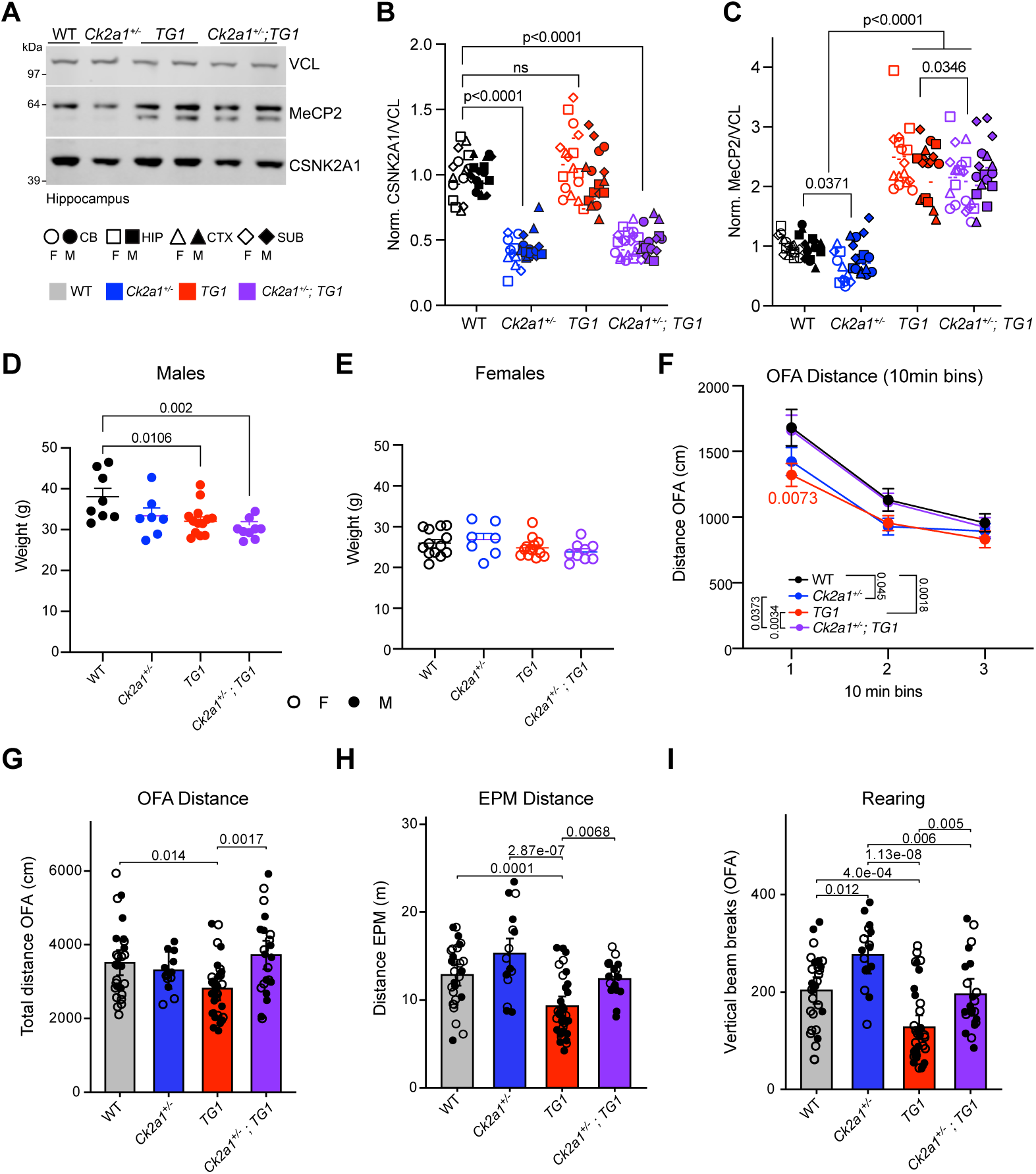
Genetic reduction of *Csnk2a1* in MDS model lowers MeCP2 levels and rescues locomotor and rearing behavior. A) Representative western blot and quantification of (B) CSNK2A1 and (C) MeCP2 protein levels relative to loading control (VCL) across brain regions of WT, *Ck2a1^+/−^*, *TG1*, and *Ck2a1^+/−^;TG1* mice (N=6-8/genotype, 3-4/sex, 4 brain regions per mouse included as nested observations). Individual datapoints are represented labelled by brain region, sex, and genotype (CB=cerebellum, HIP=hippocampus, CTX=cortex, SUB=subcortical). P-values determined using nested one-way ANOVA with Dunnett’s post-hoc multiple comparisons between WT and other genotypes and post-host Sidak’s comparison between *TG1* and *Ck2a1^+/−^;TG1* for MeCP2 and between WT and *TG1* for CSNK2A1. Weights of (D) male and (E) female mice across all genotypes. F) Total distance moved in the open field assay (OFA) measured across 10min bins or (G) total distance over 30min. H) Total distance in the elevated plus maze (EPM) assay. I) Vertical rearing behavior in OFA. N/sex for behavior: N=7F, 9M for *Ck2a1^+/−^*, N=10F, 12M for *Ck2a1^+/−^;TG1*, N=16F, 21M for *TG1*, N=19F, 10M for WT. Data presented as mean ± SEM, p-values determined using a linear or mixed model (OFA bins) with post-hoc Tukey’s multiple comparisons, see methods for details.

These mice underwent behavioral assessment, testing locomotor activity, anxiety, motor learning, grip strength, and associative memory tasks. First, we observed a sex-specific weight phenotype in male but not female mice, where *TG1* and *Ck2a1^+/−^; TG1* mice weighed significantly less than WT or *Ck2a1^+/−^* mice (**Figure 5D-E**). A known robust phenotype of both male and female *TG1* mice is hypoactivity, particularly in the open field assay (OFA) and the elevated plus maze (EPM) behavioral paradigms^14^. We observed that reduction of *Csnk2a1* in *TG1* mice significantly rescued the hypoactive phenotype in both EPM and OFA (**Figure 5F-H**). We also observed a rescue of rearing behavior in *Ck2a1^+/−^; TG1* mice, which normalized both *Ck2a1^+/−^* and the *TG1* rearing deficits (**Figure 5I**). We did not observe changes in anxiety-related phenotypes in EPM or OFA (**Supplemental Figure 5I-K**). We also tested associative memory behavior using the cued and contextual fear conditioning (FC) paradigm and found no significant rescue of *TG1* deficits (**Supplemental Figure 5L-M**). We did observe that both *TG1* and *Ck2a1^+/−^* mice had a significantly reduced response to cued FC and rescue mice were no different from WT mice (**Supplemental Figure 5L**). No rescue was observed in the *TG1* enhanced motor-learning phenotype via the rotarod paradigm and no deficits seen in grip strength (**Supplemental Figure 5N-O**). Together, these data show that reduction of *Csnk2a1* in MDS mice moderately reduces levels of MeCP2 and significantly rescues locomotor deficits and rearing behavior, with subtle effects on cued associative-memory.

### Partial rescue of MDS transcriptional disease signature with reduction of *Csnk2a1*

MeCP2 is a well-established transcriptional regulator, with the transcriptome being highly sensitive to changes in MeCP2 protein levels^34^. This is particularly true for a core set of genes that are consistently dysregulated across RTT models, known as “core MeCP2 genes”^59^. CSNK2A1 is also known to play a role in chromatin biology by phosphorylating a number of chromatin-associated proteins^60^. Given our finding that loss of *Csnk2a1* reduced levels of MeCP2 in the hippocampus, we wondered if this reduction would be reflected in downstream transcriptional changes.

Using an unbiased approach, we performed a differential gene expression (DEG) analysis on bulk RNA-sequencing (RNA-seq) of hippocampal tissue across all four genotypes in the *Ck2a1^+/−^;TG1* experimental cross using an N=6-9/genotype divided across sexes (**Figure 6A, Supplemental Table 3**). Surprisingly, we found the greatest number of DEGs in the *Ck2a1^+/−^* hippocampus (274), followed by *Ck2a1^+/−^;TG1* (205), then *TG1* (131) compared to WT samples (**Figure 6B-F**). Most upregulated and downregulated transcriptional changes were unique to each genotype, though significant overlaps were observed particularly between *Ck2a1^+/−^;TG1* and *TG1* (**Figure 6B-C**), including previous identified DEGs in *TG1* tissue like *Mastl*, *Yme1l1*, *Gdf11*, *Crhbp*, and *Cdh18* (**Figure 6E-F**)^22^. *Gdf11* has been shown in several previous studies from our lab to be directly correlated to levels of MeCP2^61^. Transcriptional changes were highly correlated between *TG1* and *Ck2a1^+/−^;TG1* samples (**Supplemental Figure 6A**), more so when examining only *TG1* DEGs (**Supplemental Figure 6B**). We performed a gene ontology (GO) enrichment analysis on *TG1*-specific, *Ck2a1^+/−^;TG1*-specific, or shared DEGs and found mostly unique GO categories for each group (**Supplemental Figure 6C**). GO categories such as locomotory behavior and adult locomotory behavior were enriched in *Ck2a1^+/−^;TG1*-specific DEGs, which is notable given the specific behavioral rescue of locomotor activity in these mice. Shared DEGs between *TG1* and *Ck2a1^+/−^;TG1* samples were enriched in GO pathways for apical plasma membrane and peptide-hormone binding (**Supplemental Figure 6C**). *Ck2a1^+/−^* upregulated DEGs were enriched for GO categories that included apical and basal parts of the cell, extracellular matrix organization and structure, cell adhesion molecule binding, and transmembrane transporter activity (**Supplemental Figure 6D**).

**Figure 6.**
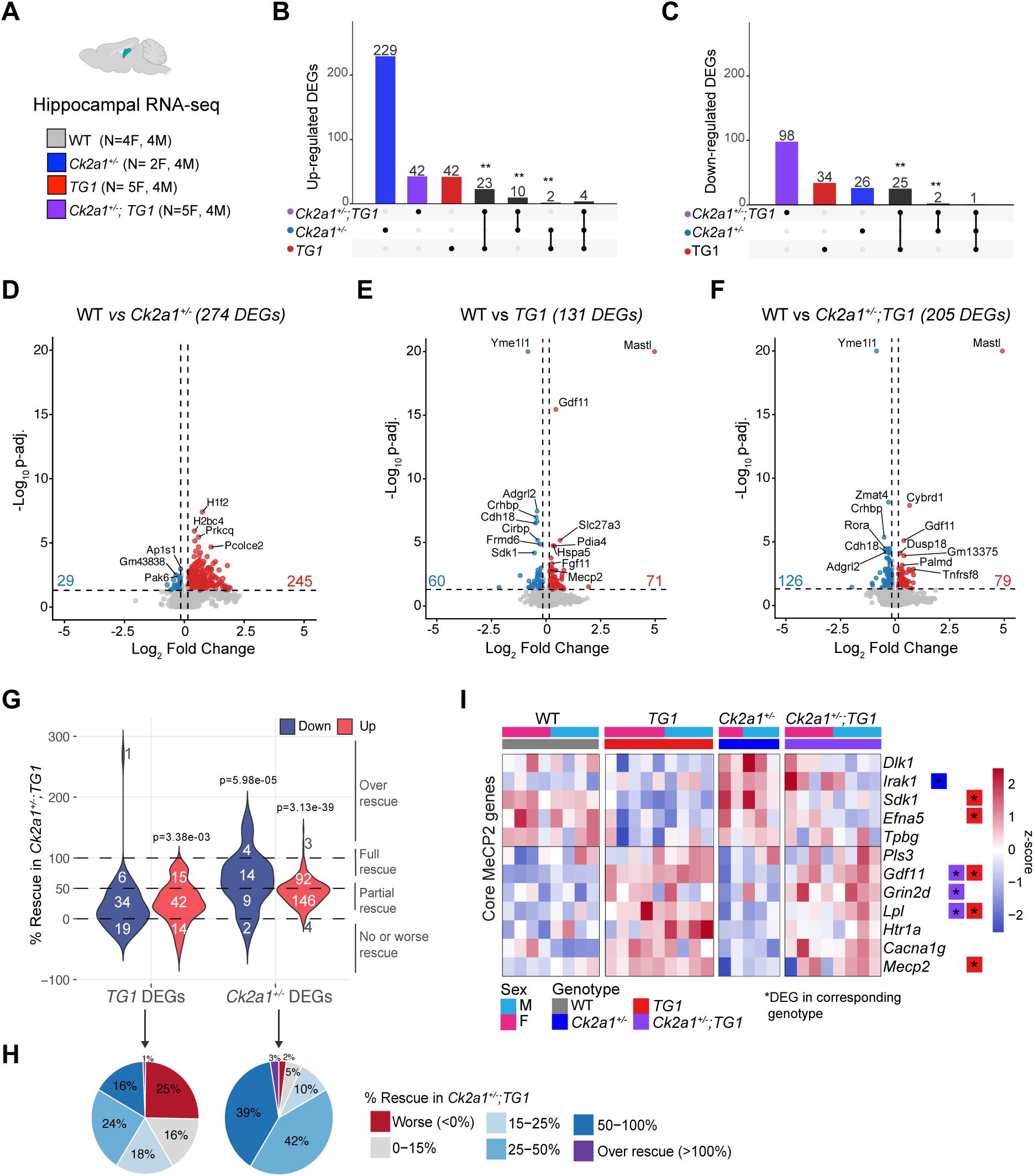
Reduction of *Csnk2a1* partially rescues MDS transcriptional signature, while MeCP2 overexpression rescues *Csnk2a1^+/−^*aberrant transcriptional activation. A) Summary of RNA-seq experiment. B-C) Upset plots showing the overlap of significant DEGs (p-adj <0.05 and log_2_FC>|.15|) across genotypes for (B) upregulated genes or (C) downregulated genes compared to WT control group ** indicate significant overlap using a hypergeometric test. D-F) Volcano plots showing the top DEGs from the RNA-seq analysis comparing (D) WT vs *Ck2a1^+/−^*, (E) WT vs *TG1*, or (F) WT vs *Ck2a1^+/−^;TG1*. G) Violin plot showing the % rescue of TG1 DEGs (left) or *Ck2a1^+/−^*DEGs in *Ck2a1^+/−^;TG1* samples, downregulated DEGS (blue) or upregulated DEGs (orange). Full or over, partial, no or worse rescue lines are indicated on the plot, with the number of DEGs in each group in white. H) Pie charts showing the percentage of *TG1* DEGs (left) or *Ck2a1^+/−^* DEGs (right) that rescued within distinct ranges in *Ck2a1^+/−^;TG1* samples. I) Heatmap of the z-score normalized expression values of core MeCP2 regulated genes across all replicates identified by genotype and sex,* indicates a significant DEG in the specified genotype.

Interestingly, we found that crossing *TG1* to *Ck2a1^+/−^* mice essentially corrected the aberrant transcriptional activation in *Ck2a1^+/−^* samples (**Figure 6D-F**). To examine the transcriptional rescue more closely, we asked how many DEGs in either *TG1* or *Ck2a1^+/−^* samples were corrected or rescued in *Ck2a1^+/−^;TG1* samples (**Figure 6G**). We were particularly interested if *TG1* DEGs moved closer to WT levels with reduction of *Csnk2a1*. Expecting subtle rescue effects, we examined the difference in log transformed normalized counts of all *TG1* DEGs in both *TG1* and *Ck2a1^+/−^;TG1* samples and found that most *Ck2a1^+/−^;TG1* gene expression levels shifted towards baseline WT levels (**Supplemental Figure 6E**). To quantify these effects, we calculated the percent rescue of each DEG from either *TG1* or *Ck2a1^+/−^* samples in *Ck2a1^+/−^;TG1* samples and found a shift towards WT levels (100%) for both up- and down-regulated DEGs (**Figure 6G**). We categorized the % rescue into distinct bins of rescue and found ∼75% of *TG1* DEGs were partially rescued in *Ck2a1^+/−^;TG1* samples, with ∼24% of DEGs rescued by 25-50%, and ∼16% of DEGs rescued between 50-100% (**Figure 6H, Supplemental Table 4**). Upregulated *TG1* DEGs were significantly shifted back to WT levels above a 15% rescue threshold in *Ck2a1^+/−^; TG1* samples. Given that modest reduction of MeCP2 protein levels in *Ck2a1^+/−^;TG1* mice, the small magnitude of rescue is what a reduction of this size would predict. Strikingly, both up and down regulated *Ck2a1^+/−^* DEGs were significantly rescued with overexpression of MeCP2, with ∼90% of DEGs partially rescued back towards WT levels, ∼40% of them rescued by greater than 50% (**Figure 6H, Supplemental Table 4**). This suggests that loss of transcriptional repression observed in *Ck2a1^+/−^* samples can be corrected with increasing the abundance of MeCP2, a known transcriptional repressor.

We next specifically examined MeCP2-sensitive transcripts, “core MeCP2 genes”^59^, across all samples and found that *TG1* samples had the expected opposite transcriptional changes as seen in *Mecp2* null mouse models, with 5 core genes reaching significance (*Sdk1*, *Efna5*, *Gdf11*, *Lpl*, and *Mecp2*) (**Figure 6I**). *Ck2a1^+/−^* samples had a significant increase in *Irak1*, while only 3 core DEGs reached significance in *Ck2a1^+/−^;TG1* samples (*Gdf11*, *Lpl*, and *Grin2d*) (**Figure 6I**). All core DEGs in *TG1* samples were partially rescued in *Ck2a1^+/−^; TG1* samples: *Sdk1* by 52%, *Mecp2* by 50%, *Efna5* by 35%, *Lpl* by 35%, and *Gdf11* by 13% (**Supplementary Table 4**). These data show that reduction of *Csnk2a1* in the MDS mouse model can modify the MDS transcriptional disease signature, particularly core genes sensitive to MeCP2 levels, underscoring that subtle changes to MeCP2 levels are reflected at the molecular level.

### MeCP2 levels are reduced in the hippocampus of OCNDS model, *Csnk2a1^K198R/+^* mice

While our discoveries have clear therapeutic implications for normalizing the pathology of MDS, they also suggest that MeCP2 dysregulation might contribute to the molecular pathogenesis of OCNDS. To test this, we used mice harboring the most frequent patient missense mutation, *Csnk2a1^K198R/+^* (K198R/+ mice). This variant is located in the p+ loop region in the activation segment domain of CSNK2A1 and occurs in ∼33% of OCNDS cases^62^ (**Figure 7A**). While the disease mechanism of this mutation is still being elucidated, previous studies have shown the K198R variant alters the substrate specificity of CK2^63^ and reduces kinase activity *in vitro*^64,65^. K198R/+ mice also display significant hippocampal-dependent learning deficits, reduced LTP, and reduced excitatory (VGLUT1+) synapses^66^. These phenotypes are similar to those observed in RTT mice that also have extensive hippocampal deficits^67–69^. Given the shared hippocampal phenotypes and our region-specific findings in *Ck2a1^+/−^; TG1* mice, we examined MeCP2 levels in the hippocampus of K198R/+ compared to littermate controls and found a significant decrease of MeCP2 by 40-50%, in addition to a ∼35% decrease in CSNK2A1 using a validated N-terminal targeting antibody^70^ (**Figure 7B-D**). This reduction was driven by post-translational mechanisms since *Mecp2* and *Csnk2a1* RNA levels were unchanged (**Figure 7E-F**). The decrease in MeCP2 was specific to the hippocampus, as we did not see any significant decreases to MeCP2 levels in other brain regions of K198R/+ mice (**Supplemental Figure 7A-O**), showing a region-specific effect of CSNK2A1 regulation of MeCP2. Interestingly, using a widely used C-terminal targeting antibody, we observed no change to CSNK2A1 levels (**Figure 7G-H**), while N-term CSNK2A1 levels were reduced to different degrees (∼10-50%) depending on brain region (**Supplemental Figure 7A-O**). Together, these data show a common OCNDS missense mutation can alter the stability of MeCP2 particularly in the hippocampus, implicating MeCP2 in the molecular pathogenesis of OCNDS. An intriguing future direction of this work will be to examine the region-specific molecular pathways regulating CSNK2A1 activity and how this changes MeCP2 stability.

**Figure 7.**
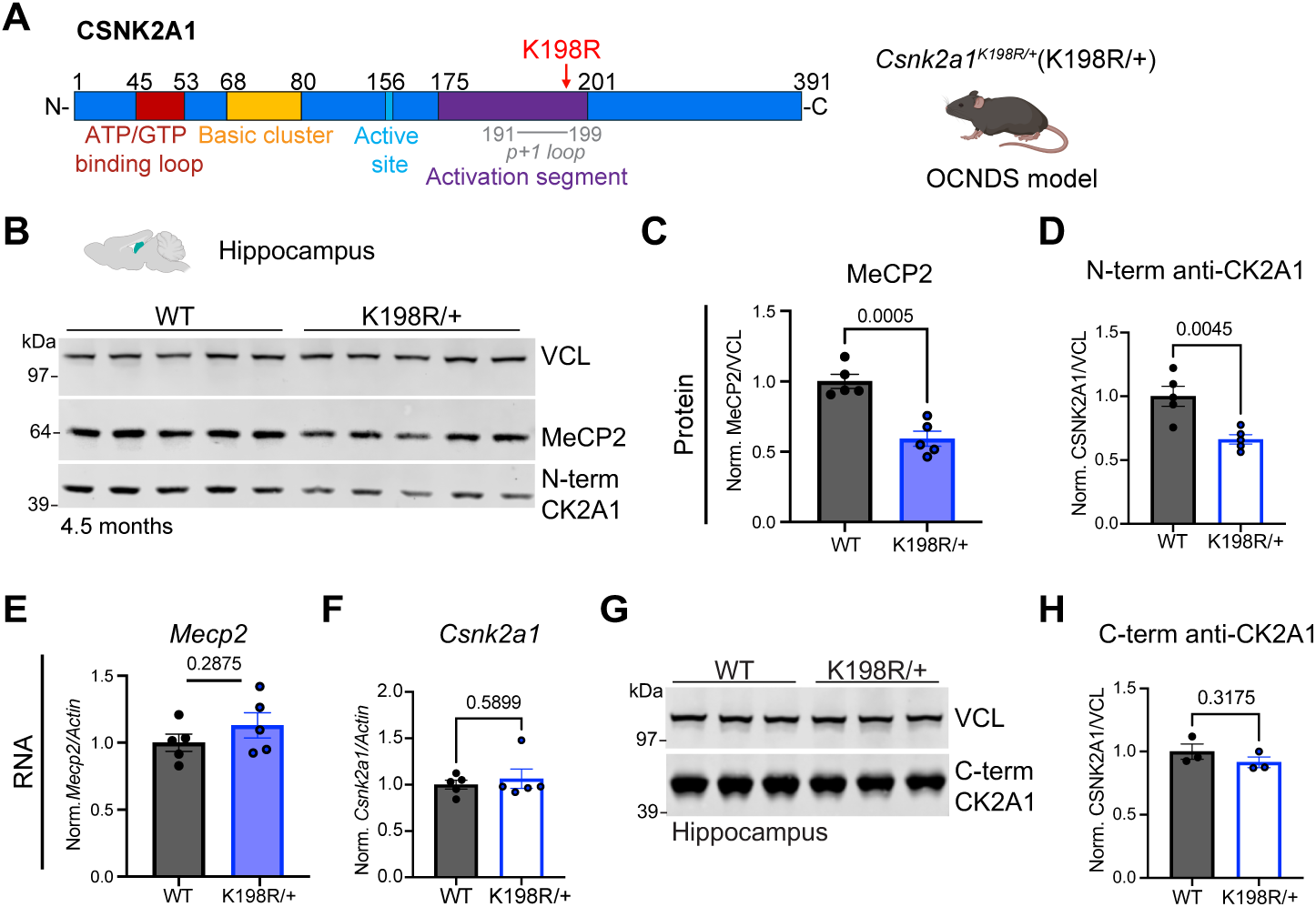
MeCP2 is reduced in the hippocampus of *Csnk2a1^K198R/+^* mice. **A**) Protein domain schematic of CSNK2A1, highlighting the most common OCNDS patient mutation, K198R, and the corresponding mouse model. **B)** Western blot and quantification of (C) MeCP2 or (D) CSNK2A1 (N-term antibody) relative to loading control (VCL) in the hippocampus of K198R/+ vs WT mice. Quantitative RT-PCR of (**E**) *Mecp2* or (**F**) *Csnk2a1* from in the same hippocampal lysates. **G-H**) Western blot and quantification for CSNK2A1 (C-term antibody) relative to loading control (VCL) in the sample hippocampal lysates of K198R/+ vs WT mice. Data presented as mean ± SEM, p-values calculated using a student’s t-test for all WT vs K198R/+ comparisons.

## DISCUSSION

Too little MeCP2 causes RTT and too much causes MDS, and in both disorders the severity tracks with how far MeCP2 levels deviate from normal. Defining the pathways that set MeCP2 protein dosage is therefore central to designing therapies that modulate MeCP2 abundance. Here, we provide the first genome-wide analysis of MeCP2 post-translational regulation by using CRISPR KO screens to identify proteins that modulate MeCP2 levels, revealing a previously unrecognized molecular pathway regulating MeCP2 stability.

Supporting the biological relevance of this approach, we found screen hits previously implicated in MeCP2 regulation, like *PPP2R1A*^24^, as well as genes more recently linked to MeCP2 biology and therapeutic interventions. For example, CDK9, a top hit stabilizing MeCP2 in our screen, is a key component of the super elongation complex (SEC), which was recently shown to interact with MeCP2 and regulate transcriptional programs involved in synaptic plasticity^71^. We also validated that STK11/LKB1, an upstream regulator of AMPK signaling^72^, stabilizes MeCP2. Consistent with this finding, activation of the AMPK pathway with Metformin was recently reported to improve the general condition of a severe RTT mouse model carrying the T158M mutation, including deficits in breathing, gait, hindlimb clasping, and tremor^73^. Together, these data highlight the biological and therapeutic potential of our functional genomic approach and provides a rich dataset for future studies aimed at understanding MeCP2 regulation.

We focused our downstream analyses on the top screen hit CSNK2A1, a highly druggable kinase encoding the catalytic alpha subunit of the well-studied CK2 holoenzyme. We show that genetic knockout, knockdown, or pharmacological inhibition of *Csnk2a1* consistently reduces MeCP2 levels both *in vitro* and *in vivo*. CK2 is a constitutively active kinase whose dysregulation has been linked to several cancers, inflammatory disorders, infection, and neurodegeneration^35,36^. In 2016, heterozygous *de novo* mutations in *CSNK2A1* were identified in individuals with a severe neurodevelopmental syndrome, OCNDS^37^, which shares overlapping clinical features with *MECP2*-related disorders, particularly RTT ^10,39^. Given these overlapping clinical phenotypes, together with our screen validation data, we prioritized investigating the molecular relationship between CSNK2A1 and MeCP2.

First, we observed that CSNK2A1 interacts with MeCP2 throughout mouse brain development, reaching peak interaction at four weeks when MeCP2 expression is highest^25,27^. This dynamic interaction coincides with critical periods of neurodevelopment, including synaptic maturation, and tracks with CSNK2A1 being an important stabilizer of MeCP2. MeCP2 is essential for proper synapse formation^74^ and phosphorylation of MeCP2, particularly at activity-dependent phosphorylation sites, plays an important role in this process^75,76^. For example, mice with serine to alanine substitutions at four activity-dependent MeCP2 phosphorylation sites (S86A, S274A, T308A, and S421A) exhibit impaired synaptic connectivity and refinement despite normal MeCP2 expression levels and the absence of RTT-like behavioral phenotypes, indicating that loss of MeCP2 phosphorylation alone is insufficient to drive RTT disease pathogenesis^76^. Likewise, we found that KO of *CSNK2A1* altered MeCP2 phosphorylation at distinct serine residues (S80, S229, S326), yet individual alanine substitutions at these residues did not change MeCP2 protein stability. Notably, mice harboring the MeCP2 S80A mutations display normal MeCP2 expression and lifespan, with only subtle motor deficits like hypoactivity^77^. Together, these data suggest that the effects of CSNK2A1 inhibition on MeCP2 stability are likely mediated either by a combination of MeCP2 phospho-sites and/or indirectly by altering phosphorylation-dependent signaling pathways or protein-protein interactions.

When we explored how CSNK2A1 modulates MeCP2 *in vivo*, we found that reducing *Csnk2a1* in *MECP2*-overexpressing MDS mice moderately lowered levels of MeCP2 and rescued deficits in locomotor activity and repetitive rearing deficits. Reduction of *Csnk2a1* also partially rescued the MDS transcriptional disease signature, including known MeCP2-sensitive genes (*Sdk1*, *Efna5*, *Gdf11*, and *Lpl*), demonstrating that even modest normalization of MeCP2 levels produces measurable molecular correction. An important finding from this study was the region-specific influence of CSNK2A1 on MeCP2 stability, where MeCP2 levels were significantly lower in the hippocampus across two distinct *Csnk2a1* mutant mouse models (*Ck2a1^+/−^* and K198R/+). Evidence of region-specific variability of MeCP2 regulation have been reported in mice with a mild, hypomorphic *Mecp2* mutation (*Mecp2-R133C)*, where MeCP2-R133C levels were significantly lower in the hippocampus compared to the cortex or cerebellum^78^. CK2 and MeCP2 are known to be important for hippocampal function and related behaviors, including synaptic plasticity such as LTP and learning and memory^66,79–81^. Since both MeCP2 and CSNK2A1 are ubiquitously expressed throughout the brain, we hypothesize that hippocampal-specific cofactors or signaling pathways contribute to the effect on MeCP2 stability, adding an important layer of complexity to the study of MeCP2 regulation.

To date, no pharmaceutical treatments are currently approved for MDS and only one FDA-approved treatment is approved for RTT, Trofinetide^82^. A goal of this study was to identify druggable pathways capable of modulating MeCP2 using a genome-wide functional genomics approach. To this end, we developed and characterized a novel inhibitor of CSNK2A1, CDD-2021, which significantly reduced endogenous levels of MeCP2 in human cells by increasing the rate of MeCP2 turnover via a CRL-dependent ubiquitin-proteasome degradation pathway. We found the magnitude of MeCP2 destabilization varied across CK2 inhibitors, with CDD-2021 producing the most robust effect on MeCP2 stability compared to TBB, without compromising cellular health at high doses like CX-4945. Importantly, CK2 also regulates many other cellular pathways, including metabolism and neuroinflammatory signaling cascades, raising the possibility that inhibition of CK2 or reduction of *Csnk2a1* might alter MeCP2 stability via indirect molecular mechanisms^35,36,83–86^. Future studies will determine whether these inhibitors, or potential analogs, provide therapeutic benefit in pre-clinical models of *MECP2*-related diseases.

Finally, our findings provide new mechanistic insights into the pathogenesis of OCNDS. First, we show a transcriptional disease signature is present in *Csnk2a1* heterozygous null mice that can be rescued by overexpression of MeCP2. Second, we find that MeCP2 levels were significantly reduced in the hippocampus of both *Ck2a1^+/−^* and K198R/+ mice. Lower levels of MeCP2 likely contribute to the hippocampal dysfunction and behavioral deficits observed in K198R/+ mice, particularly since similar hippocampal-based phenotypes have been reported in multiple RTT mouse models^69,74,79,81,87^. Moreover, the loss of N-terminal epitope signal without a change in CSNK2A1 total protein suggests that K198R alters CSNK2A1 conformation, consistent with the activation segment having extensive contacts with the N-terminal ATP/GTP-binding region^88^. These data are most consistent with reduced CSNK2A1 activity towards MeCP2^64,65^, although an altered substrate preference could produce the same result^80^. An intriguing future direction of this work will be to examine whether restoring MeCP2 levels particularly in the hippocampus of K198R/+ mice can rescue any neurological phenotypes.

In summary, we identified a network of proteins involved in regulating MeCP2 post-translationally, leading us to uncover that CSNK2A1 activity influences MeCP2 stability. Collectively, these data reveal a druggable regulatory relationship between CSNK2A1 and MeCP2, linking distinct neurodevelopmental disorders to a common a molecular nexus, and informing the molecular pathogenesis of OCNDS, *MECP2*-related disorders, and potentially other NDD-associated proteins.

## MATERIALS AND METHODS

### Generation of stable cell line

Human embryonic kidney cell T (HEK293T) were obtained from American Type Culture Collection (ATCC CRL-3216) and cultured according to manufacturer’s guidelines. Briefly, cells were grown in 37°C incubator 5% CO_2_ in complete DMEM (Fisher Scientific 10-013-CV) supplemented with 10% Fetal Bovine Serum (FBS, R&D Systems, S11150) and 1% Antibiotic-Antimycotic (Thermo Scientific 15240112). HEK293Ts were transduced with Cas9 Blasticidin Lenti Particles (Sigma-Aldrich LVCA29BST) and selected with 1ug/ml of blasticidin over several passages. Single clonal populations were obtained via FACS using a SONY SH800 sorter machine (100uM filter) on the live cell gate into individual wells of a 96-well flat bottom plate. Selected clones (23 total) that grew well were tested for Cas9 expression and functionality. Clone with highest Cas9 and gRNA KO efficiency was for downstream infection with *DSRED-IRES-MECP2-EGFP* lentiviral plasmid, cells were selected with both blasticidin (1ug/ml) and zeocin (100ug/ml) for several passages then sorted following above method. Clonal lines were selected by analyzed for quality screening metrics using flow cytometry and western blotting to test for low variations in morphology and expression of fluorescent proteins, growth rate, dynamic range of MeCP2 levels, and KO efficiency using gRNAs against both *GFP* and *MECP2*. A single Cas9 and *DSRED-IRES-MECP2-EGFP* expressing clonal line was selected for pooled CRISPR screen experiments.

### Kinome and genome-wide CRISPR libraries and viral production

LentiGuide-puro Human Kinome CRISPR Knockout Libraries (Brunello) A and B were used for the kinome screen (Addgene# 1000000082), consisting of library A gRNAs 1-4 (Addgene# 75312) and B gRNAs 5-8 (Addgene# 75313). For genome-wide library, the Human CRISPR knockout pooled Library (Brunello) was purchased as lentivirus from Addgene (73178-LV). Kinome library was amplified following suggested protocol. Briefly, each library was electroporated into STBL4 electrocompetent cells (Thermo Fisher Scientific, 11635-018) and grown for 14-16hr at 37C before plasmid purification via NucleoBond Xtra Maxi EF (Takara Bio, #740424.10). eGFP gRNAs were purchased from Addgene (#80034, #80035, #80036) and eGFP gRNAs and libraries A and B were combined at equimolar ratios before lentiviral preparation. For pooled sgRNA kinome library, lentiviral packaging was performed using TransIT-293T (Mirus, Cat #2706) in 150 mm culture dishes using a 4:3:1 ratio of lentiviral (pLenti) vector (8ug), psPAX2 (6ug), and pMD2.G (2ug) per plate. Viral media was collected at 48 and 72hrs post-transfection, filtered through 0.45um pore size filters, and frozen at −80C until needed. Viral titter was calculated using a 10-fold serial dilution of concentrated lentivirus in HEK293T cells plated at low density (∼1000) in per well in 6-well plates in triplicate and selected using 2ug/ml of puromycin until untransduced cells were completely killed (∼48hrs). Surviving colonies were visualized and counted using a filtered 0.1% crystal violet solution for at least 2 dilutions and TU/ml was the # of colonies/total volume in well X dilution factor averaged over three counts of two dilutions (∼1*10^6 TU/ml).

### Pooled CRISPR KO screens

HEK293T Cas9 and *DSRED-IRES-MECP2-EGFP* expressing cells were grown in 150 mm culture dishes and transfected at a multiplicity of infection (MOI) of 0.3 to generate a representation of 1000X for the kinome library or 500X for the whole genome library and selected with puromycin (2ug/ml) over 11-12 days while passaging cells to maintain representation. Eight experimental replicates were independently infected for the kinome screen and six for the whole-genome screen. Cells fluorescently sorted into the 10% lowest (decreased MeCP2) and 10% highest (increased MeCP2) EGFP/RFP ratio in cold FACS buffer (1X PBS, 3% FBS, and 1mM EDTA, filtered) using a SONY sorter (SH800) and control (bulk, unsorted) cells were collected at 500 or 1000X representation for genome and kinome, respectively. For each sorted replicate, cells from high, low, or bulk groups were pelleted by centrifugation at 4000 x g at 4°C for 10min, supernatant removed, flash frozen, and stored at −80C until further processing. Frozen pellets were thawed on ice and genomic DNA was extracted using the DNeasy Blood and Tissue kit (Qiagen, 69506) following manufacturer’s guidelines. gRNA sequences were amplified using two consecutive PCRs, first to amplify the gRNA sequences and second to add indices to each sample (Nextera XT indices, FC-131-2001). Final PCR was purified using both AMPureXP beads (Beckman, A63880) and sent for Pippen prep (Sage Science). DNA concentration and purity was measured using Qubit, qPCR, ad Bioanalyzer for pooling samples for Illumina HiSeq2500 using 150bp single-end reads using the Genomic ad RNA Profiling Core (GARP) at BCM. FASTQ files were analyzed using the published CRISPRcloud2 pipeline^42^. For kinome, sgRNAs against genes enriched in low or high group and depleted in the other with an FDR_pos<0.05 for either high or low groups were considered significant hits and for genome-wide, a p-value_pos <0.08 cutoff was used.

### Validation of screen hits

Single guide RNA pL-U6-puro plasmids were obtained from Invitrogen LentiArray Human CRISPR Library glycerol stocks (Cat#: A32185) following manufacturers guidelines. Dual guide RNA (pL-U6-puro) plasmids and pAAV-4-shRNA vectors were synthesized by Vector Builder. Individual lentiviruses (pLenti or pInducer) were packaged using a modified protocol described above for a smaller volume of virus in a 12-well plate, filtered, and tittered using Lenti-X GoStix Plus (Takara Bio, 631280). Purified pAAV viruses were produced through BCM Gene Vector Core and tittered using quantitative-PCR following previously published methods^89^. All guide RNA and shRNA sequences are available in Supplemental Table 5.

### Expression profile of screen hits in mouse single-cell whole brain data

Adult mouse single-cell (Allen Whole Mouse Brain, 10x v3)^45^ raw count matrices were obtained with abc_atlas_access (1.4.0). Cell types were assigned from the atlas taxonomies and cluster neurotransmitter annotation from each brain region: glutamatergic, GABAergic, mixed, cholinergic, monoaminergic, immature neuron, astrocyte, oligodendrocyte (precursors included). Remaining non-neuronal cell-types (microglia, vascular, and ependymal cells) were pooled and not displayed. Brain regions: Isocortex; OLF, olfactory areas; HIP, hippocampal region; RHP, retrohippocampal region; ENT, entorhinal area; CTXsp, cortical subplate; STR, striatum; PAL, pallidum; TH, thalamus; HY, hypothalamus; MB, midbrain; P, pons; MY, medulla; CB, cerebellum. Region × cell-type profiles are the median across donors (at least 3 mice). For each region × cell-type profile, expression of screen hits was summarized as relative expression (z-score across profiles) and absolute log_2_(CPM + 1). Unique molecular identifier counts were summed per mouse, brain region and cell type to determine raw-count pseudobulk profiles. Pseudobulk profiles were normalized to counts per million and expressed as log_2_(CPM + 1) and profiles fewer than 50 cells or 200,000 UMIs were excluded from plots. Screen hits were mapped to mouse orthologs, no statistical tests were applied to panels. Analyses used Python (3.11), with numpy (2.4.6), pandas (2.3.3), scipy (1.17.1), matplotlib (3.11.2), anndata (0.12.19). Analysis scripts were drafted with the assistance of Claude (Anthropic, Opus 5) and all code was reviewed, executed, and verified by the authors.

### Enrichment analyses of screen hits

To examine enrichment of screen hits with NDD-associated genes, we intersected genome-wide hits with inheritance-specific predictions from mantis-ml, defining risk genes as genes scoring above 90% across in each category (ASD monoallelic, DD monoallelic, DD bi-allelic, DEE monoallelic, DEE biallelic). The background used was the CRISPR library gene set targeted (19,114). One-sided Fisher’s exact test was used to calculate p-values, which were corrected across all tests Benjamini-Hochberg. Screen hits were annotated for existing drug-gene interactions by intersecting the Drug Gene Interaction Database (DGIdb v.5.0.12, 2026-06 release date). Interactions were aggregated per gene to determine the number of interacting drugs that targeted each gene with regulatory approval (Tier 1), investigational compound (Tier 2), or no recorded interaction (Tier 3). DGIdb states that these findings do not indicate the effectiveness of any drug or treatment, nor are they ranked in order of efficacy, and interactions not present do not indicate lack of effectiveness. Analyses were performed in R (4.5.2) and plotted using ggplot.

### Drug inhibitor experiments

HEK293T cells were plated in 24-well plates at 100,000 cells/well and compounds added the next day directly into the media at doses and incubation times noted in figures or figure legends. TBB (Tocris, 2275), Silmitasertib (CX-4945) (Selleck Chemicals, S2248), CDD-2021, MLN4924 (Pevonedistat, MedChemExpress, HY-70062), MG-132 10mM InSolution (Fisher Scientific, 474791), stocks were resuspended in DMSO at 10mM and diluted directly into cell culture media to specific concentrations and incubation times denoted in the figure or figure legends. Chloroquine (CQ) (Millipore sigma, C6628) stock was resuspended at 100mM in PBS. Following drug incubation, all media was aspirated and cells were flash frozen at −80C until further processing.

### Protein isolation and co-immunoprecipitation (co-IP)

Brain tissue or cultured cells were lysed in NP-40 lysis buffer (10 mM HEPES pH 7.9, 3 mM MgCl₂, 5 mM KCl, 140 mM NaCl, 0.1 mM EDTA, 0.5% NP-40/IGEPAL, 1X protease inhibitors (100X Xpert Protease Inhibitor Cocktail GenDepot P3100-020), 1X phosphatase inhibitors (100X Xpert Phosphatase Inhibitor Cocktail GenDepot P3200-020), and 25U/ml Pierce Universal Nuclease for Cell Lysis (VWR 88702). Brain tissues were lysed in 1mL of lysis buffer per 100mg of tissue. Cells were lysed In 100ul of lysis buffer per ∼1 million cells, while scrapped off plates. For brain tissue, motorized tissue grinder (handheld polytron, WPR, 47747-370) for protein/RNA extraction was used. Lysates were rotated for 30min-1hr at 4°C, then clarified by centrifugation at maximum speed for 30 min at 4°C. For co-IPs, brain tissue was dounce-homogenized in above lysis buffer. Total protein was quantified using Pierce BCA Protein Assay kit (Cat#: 23335). Co-IPs were performed using Dynabeads Protein G magnetic beads (Thermo Fisher/Invitrogen, 10004D). Beads were conjugated overnight at 4°C with either mouse anti-MeCP2 antibody (Sigma-Aldrich M6818; 4 µg per IP), mouse anti-FLAG antibody (Sigma-Aldrich, F1804; 2ug per IP), or normal mouse IgG antibody (Millipore Sigma #12-371) in 1X PBS containing 0.1% Tween-20 (PBST). Antibody-conjugated beads were washed once with PBST prior to immunoprecipitation. Lysates were pre-cleared with Dynabeads Protein G for 20 min at 4°C, beads were removed, and 10% of lysate was reserved as input. Pre-cleared lysates were rotated overnight at 4°C with msMeCP2, msFLAG, or msIgG-conjugated Dynabead beads. Beads were then washed 4X with NP-40 wash buffer lacking nuclease and eluted in lysis buffer containing NuPAGE LDS sample buffer (Thermo Fisher Scientific NP0008) and NuPAGE sample reducing agent (Thermo Scientific NP009). Input samples were prepared with the same NuPAGE LDS sample buffer and reducing agent. Samples were boiled for 5 min and either stored briefly at −20°C or immediately processed via SDS-page separation using 4-12% NuPAGE gels (Invitrogen) using MOPS-SDS Running Buffer (Boston Bioproducts, BP-178) and transferred onto nitrocellulose membrane (0.2um pore size for total protein analysis and 0.45um pore size for co-IP samples). Membranes were blocked with Li-COR Intercept (TBS) Blocking Buffer (927–60025) for 1hr at room temperature (RT) and probed with primary antibodies diluted 1:1 in Li-COR Intercept Blocking Buffer and 1X TBST. Membranes with primary antibodies incubated rocking at 4°C overnight, washed with TBST, incubated in fluorescent secondary antibodies (1:20,000) for 1hr at RT, washed, imaged on the Odyssey CLx imager (Licor), and quantified using ImageStudio software (Li-COR Biosciences).

### Antibodies

For Western blots: rbGFP (1:4,000, abcam6556), rbMeCP2 (1:5000, Cell Signaling Technology, 3456S), msCas9 (1:5,000, Cell Signaling Technology, 14697S 7A9-3A3), msGAPDH (1:25,000, Advanced ImmunoChemical, 2-RGM2 Clone 6C5), rbCDK9 (1:1000, Cell Signaling Technology, 2316T), rbBRD4 (1:1000, ThermoFisher Scientific, 301-985A-M), rbLKB1 (also known as STK11, 1:1000, Cell Signaling Technology, 3050S), rbTAF1 (1:1000, Protein tech, 20260-1-AP), and rbCSNK2A1 (1:5000, Protein-tech, 66627), msVCL (1:5000, Sigma-Aldrich V9131), rb-phospho-MeCP2(Ser-80) (1:10,000, EMC Bioscience, MP4601), rbHA (1:10,000, Cell Signaling Technology 3724S), msFLAG (:10,000, Sigma-Aldrich, F1804). For CSNK2A1, we used two additional antibodies, rbCSNK2A1 (1:5000, abcam76040, for N-terminal targeting within the first 100 amino acids of CK2A1) and rbCSNK2A1 (1:5000, Bethyl Ab, A300-198A, for C-term).

### qPCR analysis

RNA was isolated from cultured cells or mouse brain tissue using RNeasy Mini kit (Qiagen, 74106). DNA was removed using Invitrogen DNAse I treatment (Thermo Fisher Scientific, 18068015) and cDNA synthesized using Primescript RT Master mix (Takara Bio, RR036A) following manufacturers guidelines. qPCR was performed using PowerUP SYBR Master Mix (Thermo Fisher Scientific, A25778) and gene-specific primers spanning exon junctions (designed using Primer3 input) on Bio-Rad CFX96 real-time PCR machine. Relate transcript abundance was calculated using the ddCT method relate to a housekeeping gene (*ACTIN* or *Actin* for human or mouse samples). Primer sequencing can be found in Supplemental Table 5.

### Synthesis of CDD-2021

CDD-2021 was synthesized at the Center for Disease Discovery (CDD) at Baylor College of Medicine following previously published methods^50–53^. Extended experimental procedures for synthesis and NMR data are in supplemental general chemistry methods. Briefly, all starting materials and reagents were purchased from commercial sources and used without further purification. Solvents were purchased as either anhydrous grade products in sealed containers or reagent grade and used as received. All reactions were carried out in dry glassware under a nitrogen atmosphere using standard disposable or gastight syringes, disposable or stainless-steel needles, and septa. Stirring was achieved with magnetic stir bars. Flash column chromatography was performed with SiO2 (230-400 mesh) or by using an automated chromatography instrument with an appropriately sized column. Thin layer chromatography was performed on silica gel 60F254 plates (E. Merck). Non-UV active compounds were visualized on TLC using one of the following stains: KMnO4, ninhydrin, p- anisaldehyde. 1H and 13C NMR spectra were recorded on an instrument operating at either 600MHz, and 151MHz respectively. LCMS data were collected using an HPLC instrument coupled to a low-resolution mass spectrometer with single quadrupole ionization operating in either positive or negative ion mode. The analytical method utilized a C18 column (2.1 × 50 mm, 1.8 µm) eluting with a linear gradient of 95%/5% water/CH3CN (modified with 0.05% formic acid; T = 0 min flow = 0.35 mL/min) to 95%/5% CH3CN/water (T = 3.5 min flow = 0.5 mL/min) then 95%/5% CH3CN/water to T = 5min (0.5 mL/min). Peak detection was done at 254 nm and 230 nm for UV active compounds. High-resolution mass spectrometry (HRMS) spectra were obtained on a Thermo Scientific Q Exactive hybrid quadrupole-Orbitrap mass spectrometer equipped with a HESI source and using lock masses for correction. Samples were introduced into the HRMS via reversed phase HPLC on an Accucore Vanquish C18+ column (2.1 × 100 mm, 1.5 µm) eluting with a linear gradient of 95%/5% water/acetonitrile (modified with 0.1% formic acid) to 10%/90% water/acetonitrile over 8 min.

### LC-MS/MS Analysis

The protein complex was eluted from the beads in NuPAGE™ LDS Sample Buffer and resolved on 10% Bis-Tris NuPAGE gel (Life Technologies), visualized with Coomassie Brilliant blue-stain. The target protein band region from 60-160kDa was excised and in-gel digested with trypsin and LysC protease mixture. The peptides were concentrated in a speed vac and dissolved in 12µl of 5% methanol containing 0.1% formic acid loading solution. The LC-MS/MS analysis was carried out on a Vanquish Neo UHPLC system (Thermo Fisher Scientific, San Jose, CA) coupled to Orbitrap Eclipse (Thermo Fisher Scientific, San Jose, CA) mass spectrometer. The MS data was acquired for 110min gradient time on a 20cm in-house packed column in a data dependent acquisition mode. The MS1 was done in Orbitrap (120000 resolution, scan range 375-1500m/z, 50ms Injection time) followed by MS2 in Ion Trap with CID fragmentation. The MS raw data was searched in Proteome Discoverer software (v1.4, Thermo Scientific, San Jose, CA) with Mascot algorithm (v2.4, Matrix Science, PMID 10612281) against the mus musculus or homo sapiens protein database downloaded from NCBI Refseq (2021_12_23). Precursor and fragment mass tolerance were set to 20ppm and 0.5Da respectively. Database searches allowed a maximum of 2 missed cleavages with trypsin/P enzyme specificity. Dynamic modifications included oxidation (M), protein N-terminal acetylation, deamidation (N/Q), phosphorylation (S/T/Y), methylation (R) and carbamidomethylation (C). Peptide identifications were validated at a 5% false discovery rate (FDR) using Percolator node (PMID 17952086). The modified peptide spectra were manually curated for correct site assignment. Area under the curve for peptide quantification was obtained using Skyline (v23.1, MacCoss Lab, University of Washington, Seattle, WA) by importing result files from Proteome Discoverer. The unmodified peptides were used for normalization across samples.

### ADP-Glo kinase assay and recombinant protein purification

ADP-Glo kinase assay + CK2 alpha 1 Kinase Enzyme System (Promega, V4483) was used following manufacturers guidelines. Recombinant MeCP2 was made using a pET-GST-MECP2 (human e2) plasmid from Vector Builder. BL21(DE3) competent cells from NEB (C2527) were used for transformation and single colonies were grown in 50ml LB broth, supplemented with 50ug/ml ampicillin, shaking at 250 rpm overnight. Culture was induced by adding 0.1mM IPTG and incubated for 4h at 37C with shaking. Cells were harvested with centrifugation and lysed in cold PBS via sonication. Cellular debris was removed by centrifugation (15,000 x g for 15min at 4C) and supernatant collected for purification using glutathione-sepharose beads. Briefly, beads were washed with cold PBS and supernatant incubated overnight at 4C. Beads were washed 4X with cold PBS and eluted with 200ul of freshly prepared elution buffer (100nM Tris pH 8.0, 120mM NaCl, 10mM reduced l-glutathione) rotating overnight at 4C. Beads were spun at 5000 xg for 1min and supernatant removed, aliquoted, and frozen at −80C. Pre- and post-induction, supernatant, pellet, and post-purified protein samples were analyzed via SDS-page gels followed by Coomassie blue staining and western blot methods probing for total protein and MeCP2 antibody (rbMeCP2, Cell Signaling 3456S). ADP-Glo assay was performed in quadruplicate using the 96-well format following manufacturer’s protocol +/−CDD-2021 (1uM) or DMSO with 100ng of CSNK2A1 and 1ug of MeCP2 or casein in 25ul reaction volume with 10uM ATP, and 60min RT incubation. Bioluminescent signal was detected on a GloMax Discover instrument (Promega). Quantification is luminescence minus matched no-enzyme background for each substrate and treatment relative to average casein + DMSO signal.

### Ubiquitination assay

HEK293T cells were seeded in 12-well plates and transfected using Mirus at ∼70% confluency with pcDNA3.1-HA-Ub and pcDNA3.1-Flag-MECP2-e2. Cells were incubated with CDD-2021 (250nM) or DMSO control for 48hrs. MG-132 (10uM) or CQ (40uM) or DMSO control was directly added to media for 8-10hrs prior to harvest by removing media and freezing cells at −80C until downstream processing. Protein was isolated and immunoprecipitation using msFLAG antibody was performed as described above.

### Cloning

Doxycycline (DOX) inducible plasmid pInducer20^56^ with puromycin resistance was provided by Dr. Tiemo Klisch and MECP2-e2 overexpression construct was cloned into this vector using Gibson assembly reaction (NEB E5510S). pInducer20 constructs using Flag-MECP2-e2 with serine to alanine mutations (S80A, S229A, S326A) were cloned using NEB site directed mutagenesis kit followed by Gibson assemble reaction. CSNK2A1 overexpression construct was ordered from Horizon Discovery (OHS5893-202494275) and cloned into the pLenti-EF1a (Addgene #85132) using Gibson reaction.

### DOX-inducible experiments

Stable HEK293T cells lines were created for each pInducer20 plasmid using lentivirus infection followed by selection with 1ug/ml puromycin until non-infected cells were completely killed. Stable cell lines were passaged 2-3X before plating in 12-well dish at ∼100,000 cells/well corresponding to each timepoint (72hr, 48hr, 24hr, 0hr without DOX). DOX (Sigma-Aldrich, D9891) stocks were made in PBS at 1mg/ml and added to cells 24hrs after plating at 100ng/ml except for no-DOX control wells. For TBB and CDD-2021 experiments, drugs were added along with DOX at 10uM and incubated for 48hrs. DMSO was used as vehicle control. DOX was removed at each timepoint and fresh media +/−DOX +/−inhibitors was replaced at each timepoint. All plates were collected at the same timepoint to control for differences in confluency, media was completely removed, and plates were frozen at −80C until protein isolation processing.

### Animals

All mouse work was approved by the Institutional Animal Care and Use Committee (IACUC) at Baylor College of Medicine animal protocol number AN-1013. *Csnk2a1^flox/flox^* mice^57^ received from Dr. Etty Benveniste from UAB. *Sox2-Cre* (B6.Cg-Edil3^tg(Sox2-Cre)1Amc^/J) female mice were purchased from the Jackson Laboratory (strain # 008454) and crossed to male *Csnk2a1^flox/flox^* mice to generate constitutive Csnk2a1 heterozygous mice (*Csnk2a1^+/−^*). *MECP2-TG1* mice were previously described and available through the Jackson Laboratory (strain # 008679). *Csnk2a1^K198R/+^* mice purchased from the Jackson Laboratory (MMRRC strain #67399-JAX). All genotyping was performed according to published guidelines or protocols available through Jackson Laboratory. Primer sequences provided in Supplemental Table 5. For AAV ICV injections and developmental timepoints, E16.5 timed pregnant Swiss Webster (CFW) mice were ordered through Charles River (strain code: 24). Behavioral testing was performed on both male and female mice of four genotypes (WT, *Ck2a1^+/−^, MECP2*-TG1, and *Ck2a1^+/−^*;*MECP2*-TG1) maintained on a C57BL6/J background and generated from crossing WT females to male *Ck2a1^+/−^; MECP2-TG1* mice across 3 cohorts. Animals were group-housed under a [14:10] light– dark cycle and tested during the light phase, with food and water available *ad libitum*. All mice were between 25-35 weeks of age at the onset of behavioral battery. All experimenters were blind to genotype throughout testing and scoring.

### ICV injections of AAV virus

ICV injections were performed on P0 pups as previously described ^90^. Briefly, AAV8 viruses with shRNAs targeting *Csnk2a1* or non-targeting (NT) were diluted to 3.5e10 viral units/ul in sterile PBS and 0.05% filtered trypan blue solution. P0 pups were cryo-anesthetized until immobile and 2ul of diluted AAV virus (7e10 vial units per hemisphere) were bilaterally injected into the lateral ventricle. Trypan blue was used to visually note successful injections by spreading throughout the ventricle. Pups from the same litter were injected with either shCsnk2a1 or shNT (∼6-7 pups injected per AAV shRNA). Mice were harvested at 8 weeks post-injection for protein and RNA analyses on MeCP2 or at 4 weeks to check viral spread by immunofluorescent staining.

### Brain tissue preparation for protein and RNA

Mice were sedated using isoflurane inhalation before the brain was removed and brain regions (olfactory bulb, cortex, cerebellum, hippocampus, striatum, and subcortical) dissected on ice-cold surface and immediately frozen on dry ice and stored at −80C until further processing. For AAV ICV injection, cortex and dorsal hippocampus was harvested and lysed together for protein and RNA analyses. Frozen samples were homogenized in PEPI buffer (1X PBS, 5mM EDTA, protease inhibitor cocktail, and phosphatase inhibitor cocktail) using 1ml of buffer per ∼100mg of tissue. Aliquot of the homogenate was diluted 1:1 in NP-40 lysis buffer (see above) and rotated at 4C for 30min followed by centrifugation at 15,000 rpm for 15min. RNA was extracted from the same homogenate using Trizol following manufacturers guidelines in RNeasy mini kit (Qiagen).

### Brain tissue fixation and sectioning

Adult mice were given a subcutaneous injection of analgesia (Buprenorphine, 0.5mg/kg) 30 minutes prior to anesthesia (Rodent Combo III, 1.5ml/kg) via intraperitoneal injection. Mice were perfused with ice-cold PBS followed by 4% PFA. Brains were removed and placed in 4% PFA overnight. Brain samples were then washed with ice-cold PBS and stored in 30% sucrose with 0.01% sodium azide. Left hemisphere of brain was sectioned along the sagittal plane at 40um thickness on a Leica SM2010 R sliding microtome and free-floating sections were stored in PBS + 0.01% sodium azide.

### Immunofluorescent staining and imaging

Brain sections were washed with PBS (3X) and incubated in permeabilization buffer (0.3% Triton-X-100 in PBS) for 5min at room temperature. Sections were placed in blocking buffer (0.3% Triton-X, 5% normal goat serum, 5% normal donkey serum, and 1% BSA) for 30min at room temperature and then incubated with primary antibodies in blocking buffer overnight at 4C. Sections were washed with PBS (3X) then incubated with secondary antibodies in blocking buffer for 1hr at room temperature. Primary antibodies used: mouse anti-NeuN (1:1000, Millipore Sigma, MAB377), rabbit anti-MeCP2 (1:1000, Cell signaling 3456S), and chicken anti-GFP (1:1000, abcam, ab13970). Secondary antibodies were all used at 1:1000 dilutions: donkey anti-mouse 647 (Jackson ImmunoResearch 715-605-150), donkey anti-rabbit 555 (Invitrogen a31572), and goat anti-chicken 488(Invitrogen, A-11039). After final washes with PBS, sections were mounted onto slides and coverslips were placed using Prolong gold antifade mountant (ThermoFisher Scientific P36934). Slides were allowed to dry overnight and then z-stack, representative images were taken using a Zeiss LSM880 confocal microscope.

### Dual-color fluorescence RNA in situ hybridization (ISH)

ISH was performed by the RNA In Situ Hybridization Core at Baylor College of Medicine using an automated platform^91^ and previously described methods^92^. Briefly, adult mouse sagittal sections from fresh frozen brain tissue embedded in OCT and frozen on dry ice were sectioned using a Leica CM3050 cryostat at 25um thickness. Digoxigenin (DIF) and fluorescein (FITC)-labeled mRNA probes for *Csnk2a1* and *Mecp2* (from Allen Brain Atlas sequences) were generated using PCR on mouse cDNA as template and DIG or FITC added by RNA labeling kit from Roch. Slides were imaged on a Leica SP8X confocal microscope using 20X and 40X objectives.

### Behavior

All assays were conducted during the light phase of the cycle, in a dedicated behavioral testing suite. Background white noise was maintained at 60 dB using a white noise generator except where noted. Mice were habituated to the testing environment for 30 min prior to each assay, and apparatus surfaces were cleaned with 35% Isopropanol between animals. Assays were administered in the following order: elevated plus maze, open field, accelerating rotarod, grip strength, and fear conditioning with a minimum of 24 hours separating consecutive assays.

### Elevated Plus Maze

Room illumination was adjusted to 150 lux and background noise to 60 dB. Following habituation, each mouse was introduced to the central zone of an elevated plus maze comprising two open arms (length: 25 cm; width: 8 cm) and two closed arms with walls on either side (length: 25 cm; width: 8 cm; wall height: 15 cm) oriented to face an open arm, and permitted to explore freely for 10 min. Time spent in each arm was measured using ANY-maze (Stoelting Inc.). Parameters extracted included time spent within each arm, frequency of zone-to-zone transitions and cumulative distance traveled.

### Open Field Assay

Room illumination was adjusted to 150 lux and background noise to 60 dB. Following habituation, mice were placed into a clear plexiglass arena (40 × 40 × 30 cm) and monitored for 30 min via infrared beam interruption. Horizontal and vertical beam breaks (activity counts), total distance traveled, and both entries into and time spent within a centrally defined 10 × 10 cm zone were acquired and quantified with AccuScan Fusion software (Omnitech Electronics Inc.).

### Accelerating rotarod

Testing was performed under ambient room illumination with background white noise at 60 dB. Following habituation, mice were placed on an accelerating rotarod (Ugo Basile) programmed to ramp from 5 to 40 r.p.m. over 10 min, following prior reports in which *MECP2*-TG1 animals outperform wild-type littermates on this task and would otherwise be censored at the shorter limit (Collins et al., 2004). Animals completed four trials per day across four consecutive days, with a minimum inter-trial interval of 30 min. Latency to fall from the rotating rod was recorded for each trial; passive rotation was scored as a fall.

### Grip strength

Forelimb and all-limb grip force were measured in three consecutive pulls using Coulbourn Instruments. Plots show the mean of 3 replicates per animal.

### Fear Conditioning

Mice were habituated in a room adjacent to the testing suite. On the training day, animals were transferred to the testing room (ambient illumination, no background noise) and placed individually into conditioning chambers fitted with a shock-delivering grid floor (ActiMetrics) each enclosed within a sound-attenuating cubicle equipped with a digital camera, loudspeaker, and house light. The training protocol began with a 2-min baseline period free of tone or shock, followed by a 30-s tone (85 dB) followed by a 2-s foot shock (0.8 mA); after a further 2-min interval, this tone–shock pairing was delivered a second time. Animals were returned to their home cages upon completion. Testing was conducted 24 hr later and comprised contextual and cued components. For the contextual test, mice were returned to the original conditioning chamber for 5 min in the absence of tone or shock. One hour later, cued testing was performed in a novel context (white plexiglass inserts placed on the floor and sides, vanilla extract for new odor, testing room under dim red light, mice brought to the testing room in transfer cages that contained paper towels instead of corn cobb bedding) over 6 min: 3 min without stimuli, followed by 3 min of tone presentation alone (85 dB). Behavior was video-recorded and scored automatically, with freezing defined as the cessation of all movement apart from respiration, in FreezeFrame (ActiMetrics) using a minimum bout duration of 1s and a motion threshold of 10.

### Statistical analysis

Statistical analyses for protein and RNA data were performed in GraphPad Prism 10 and exact statistical tests with multiple comparisons are described in each figure legend. Behavioral analyses were performed in R 4.5.2 using lme4 (2.0-1), lmerTest (3.2-1), and emmeans (2.0.3). Single-timepoint measures (elevated plus maze, open field, contextual and cued freezing, grip strength) were analyzed using a linear model with genotype as factor of interest and covariates being sex, age, and cohort. Adjusted p-values determined using Tukey’s post-hoc multiple comparisons tests. Outliers were detected using 1.5X interquartile-range criteria and excluded prior to modeling. For repeated-measures endpoints (open field distance across 10-min bins and rotarod latency across 4 days), a linear mixed model with genotype, time, and interaction as fixed effects with same covariates as above. Repeated-measures data were analyzed without outlier exclusion, since mixed model accounts for within-animal variability. Data are presented as mean ± SEM, and significance was defined as p< 0.05. Exact N values, test statistics, and p*-*values for each comparison are reported in figure legends.

### RNA-seq: RNA isolation, library preparation, and sequencing

Total RNA was isolated using the QIAzol lysis reagent (Cat # 79306) and purified using the RNeasy kit (Cat #74106) following manufacturer instructions. Total RNA (2ug) was sent to Genewiz (from Azenta Life Sciences) for library preparation using polyA selection for mRNA species, followed by strand-specific, paired-end sequencing on an Illumina sequencing platform NovaSeq X Plus (2 × 150 bp). Raw FASTQ files were used for downstream analysis.

### Read processing and alignment

Raw reads were assessed using FastQC (v0.12.1) and summarized with MultiQC (v1.15). Adapter sequences and low-quality bases were trimmed using fastp (v0.23.4) with automatic adapter detection, per-base quality trimming, and length filtering. Post-trimming quality was re-evaluated using FastQC and MultiQC to confirm read quality and adapter removal. Reads were aligned to the Mus musculus reference genome (GRCm39) using STAR (v2.7.11a). Genome indices were generated using GENCODE vM36 gene annotations with splice junction overhang set to read length minus one (sjdbOverhang = 149 for 150 bp reads). Alignment was performed using sorted BAM output mode (--outSAMtype BAM SortedByCoordinate) and strand specificity was determined using RSeQC prior to final alignment. Gene-level counts were obtained from STAR ReadsPerGene.out.tab output using the strand-appropriate column. Alignment metrics, including uniquely mapped reads, multimapping percentage, and unmapped reads, were extracted from STAR Log.final.out files and summarized across samples.

### Count Matrix processing

Gene-level count matrices were compiled across samples and imported into DESeq2 (v1.50.2.) in R (v4.5.2). Lowly expressed genes were filtered prior to modeling. Count normalization was performed using DESeq2’s median-of-ratios method. Batch effects were assessed and corrected where necessary using design formulas incorporating batch covariates. Principal component analysis (PCA) was performed on variance-stabilized counts to evaluate sample clustering and identify potential outliers. Two samples were removed upon outlier analysis (1M *Ck2a1^+/−^*, 1F *Ck2a1^+/−^; TG1* sample).

### Differential expression analysis

Differential gene expression (DEG) analyses were conducted using DESeq2 with genotype as the primary contrast variable. Statistical significance was determined using Wald tests with Benjamini–Hochberg correction for multiple testing. Genes with adjusted p < 0.05 and absolute log2 fold change ≥ 0.15 were considered differentially expressed. Gene annotations were mapped using GENCODE vM36 reference GTF files via the Bioconductor package rtracklayer. Sensitivity analyses were performed by repeating differential expression analysis with and without PCA-discordant samples to confirm robustness of identified gene expression changes.

### Transcriptional rescue analysis

To quantify the percent rescue of TG1 or CK2A1 disease-associated changes in Ck2a1; TG1 mice, we calculated the percent rescue of significant TG1 or CK2A1 DEGs using mean VST-normalized expression values for each gene using and the following: % rescue= 100* ((TG1-CK2A1;TG1 or CK2A1)/(TG1 or CK2A1-WT)). A value <0 means worsened, 0 means no rescue, 0-100 means partial rescue, 100 means full rescue, and >100 is an overcorrection. Significance of % rescue for each group was determined using a one-sample Wilcoxon signed-rank test with multiple testing using the Holm correction, testing whether the median % rescue exceeds 15%. Log_2_ Fold correlation analyses were performed on values from TG1 or Ck2a1; TG1 samples and DEGs specific to each genotype or shared in both genotypes are highlighted in each plot.

### Gene ontology analysis

Gene ontology analyses (GO) were performed using Entrez gene identifiers from the mouse annotation database org.Mm.eg.db and performed separately for significant DEGs that were either upregulated or downregulated or specific or shared between genotypes using clusterProfiler. All DEGs mapped to Entrez gene identifiers were used as background. Enrichment was evaluated across Biological Process (BP), Molecular Function (MF), and Cellular Component (CC) GO ontologies, with multiple-testing correction performed using Benjamini-Hochberg. GO terms with adj. p<0.05 were significant and results were visualized as gene ratio, number of genes contributing to each term, and adj p-value.

### Data visualization

Volcano plots, heatmaps, violin plots, and PCA visualizations were generated in R using ggplot2, pheatmap, and DESeq2 visualization functions.

### Data availability

Datasets generated from this study will be made publicly available upon the date of this publication. All code will be deposited at Zenodo upon the date of this publication. Any additional information required to reanalyze the data reported in this paper is available from the lead contact upon request.

## Supporting information

Supplemental chemistry methods

## ACKNOWLEDGEMENTS

We would like to thank all the support from the BCM cores including the GARP, Gene Vector, RNA ISH, and Proteomics Cores. We would also like to thank Dr. Hari Yalmanchili and his lab for developing the bulk RNA-seq data analysis pipeline.

## AUTHOR CONTRIBUTIONS

Conceptualization: AGA, DY, HYZ

Methodology: AGA, SC, RMS, EHC, JPR, JK, MZ, SV

Investigation: AGA, SC, RMS, JK, MZ, SV

Formal Analysis: AGA, SC

Visualization: AGA, SC

Supervision: DY, HYZ

Writing—original draft: AGA

Writing—review & editing: AGA, SC, SV, RMS, JK, DY, HYZ

Funding Acquisition: DY, HYZ

Project administration: HYZ

## DATA AND CODE AVAILABILITY

Datasets generated and code from this study will be made publicly available upon the date of this publication. Any additional information required to reanalyze the data reported in this paper is available from the lead contact upon request.

## SUPPLEMENTAL FIGURES

**Supplemental Figure 1.**
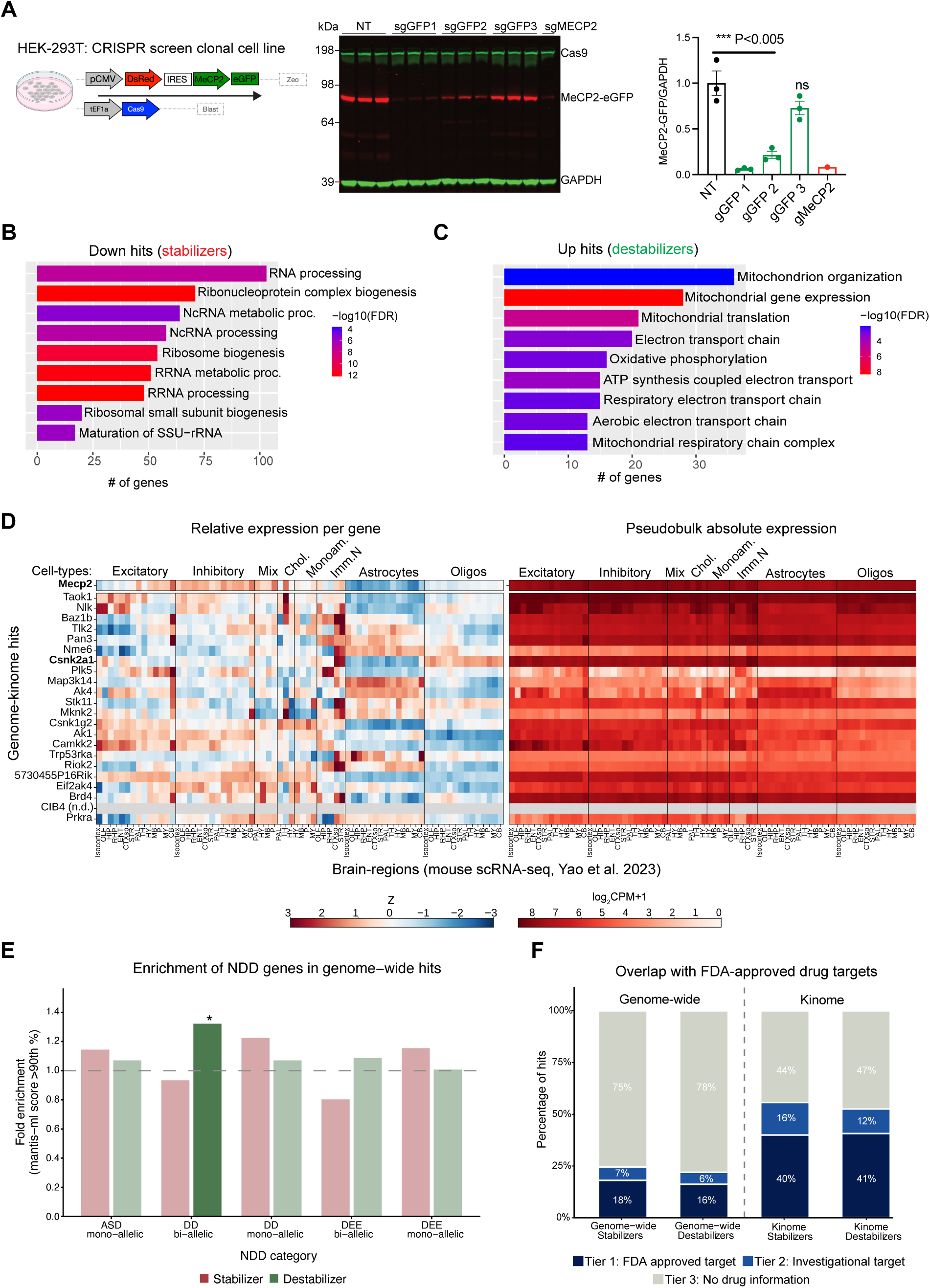
Characterization of human HEK-293T-Cas9 bicistronic MECP2 reporter cell line and CRISPR screen hits. A) Western blot and quantification of MeCP2 levels relative to loading control (GAPDH) using sgRNAs targeting *GFP* (1-3) and an sgRNA targeting *MECP2* in the CRISPR screen cell line, ***p<0.005 using a one-way ANOVA with post-hoc Dunnett’s multiple comparisons. B-C) GO categories of genome wide screen hits that either (B) down-regulated MeCP2 (stabilizers) and (C) up-regulated MeCP2 (destabilizers). D) Cell-type specific expression of genome-kinome overlapping hits across all brain regions from the Allen mouse brain scRNA-seq atlas with *Mecp2* expression shown as a reference. Left heatmaps show relative enrichment of each gene across brain regions and cell-types, while the right heatmaps shows absolute expression levels of each gene (n.d= hits not detected in single-cell data). E) Fold enrichment (observed/expected overlap) of stabilizer or destabilizer screen hits with predicted NDD genes (scoring above 90^th^ percentile across five inheritance-specific mantis-ml models) (*43*). F) Percentage of genome-wide and kinome screen hits separated by direction and stacked by approved drugs (Tier 1), investigational compounds (Tier 2), or no recorded drug-gene interaction (Tier 3) from DGIdb.

**Supplemental Figure 2.**
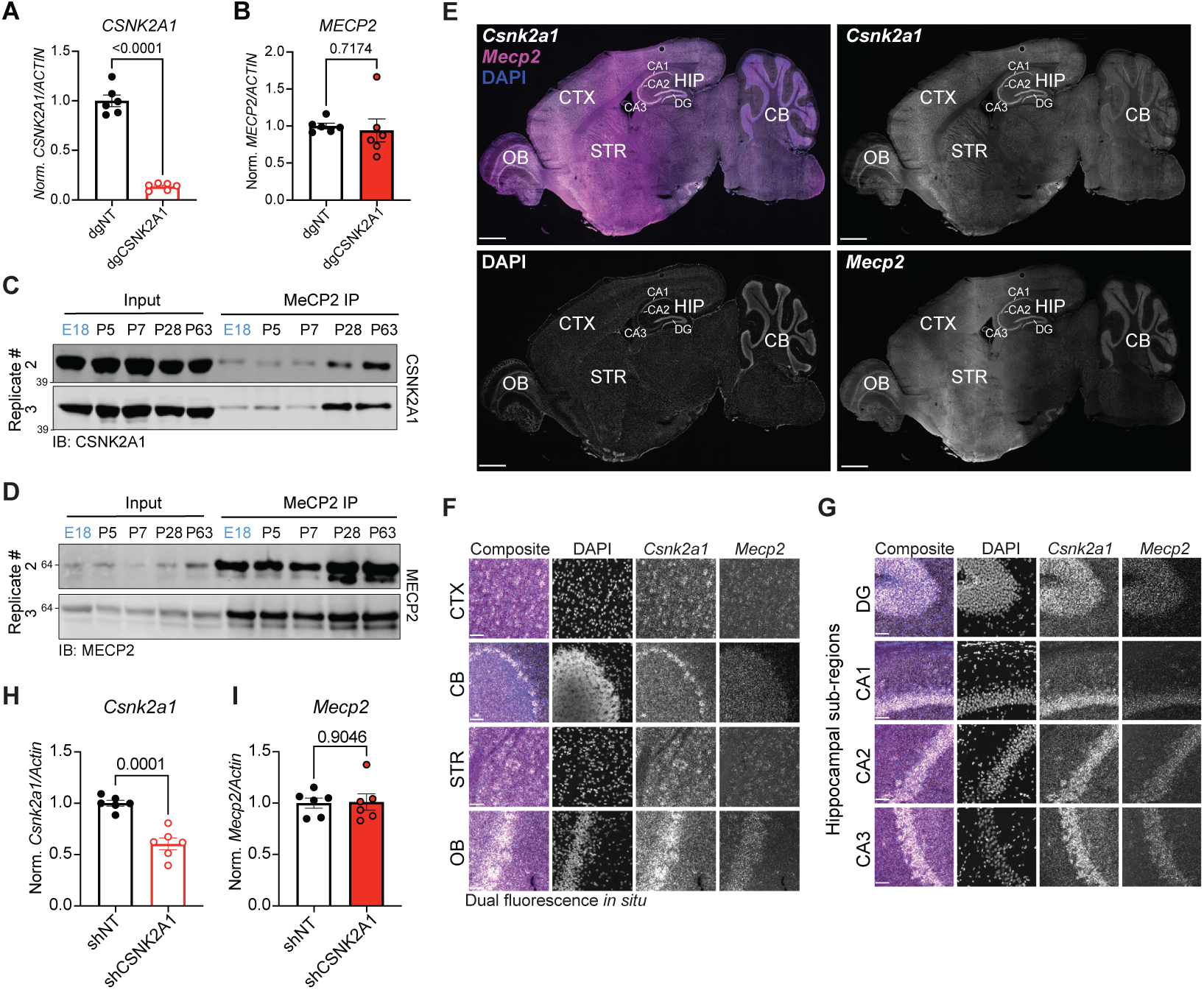
CSNK2A1 and MeCP2 binding over development and *in situ* expression in adult mouse brain. A-B) qRT-PCR of (A) *CSNK2A1* and (B) *MECP2* relative to *ACTIN* (housekeeping gene) in HEK293T-Cas9 cells infected with either dgCK2A1 (KO) or dgNT (control). C-D) Replicates (2 and 3) of IP for MeCP2 at embryonic (E18) and post-natal timepoints (P5, P7, P28, and P63) blotting for (C) CSNK2A1 or (D) MeCP2. (E-G) Dual *in situ* probes for *Mecp2* and *Csnk2a1* and stained with DAPI in a representative adult sagittal brain section at 10X (E) or 40X oil (F-G). OB= olfactory bulb, CTX= cortex, STR= striatum, HIP= hippocampus, CB= Cerebellum. Cornu Ammonis (CA) 1-3 and the dentate gyrus (DG) are subregions of the hippocampus. Scale bar= 1mm (E), 500um (F-G). H-I) qRT-PCR from AAV-*shCsnk2a1* or *shNT* P0 mouse ICV injections quantifying (H) *Csnk2a1* and (I) *Mecp2* relative to *Actin*. Data presented as mean ± SEM, p-values using a Student’s t-test.

**Supplemental Figure 3.**
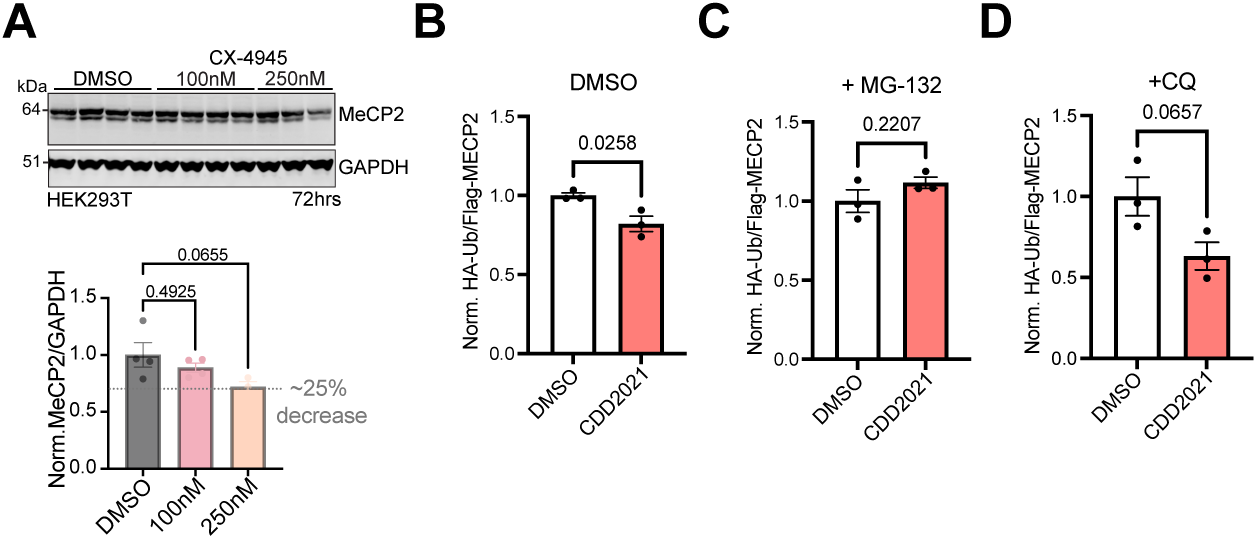
Testing CX-4945 and CDD-2021 dependent MeCP2 degradation pathways. A) Western blot and quantification of MeCP2 levels relative to loading control (GAPDH) in HEK293T cells incubated with CX-4945 at low doses for 72hrs. Data presented as mean ± SEM, p-values determined using a one-way ANOVA with post-hoc Dunnett’s multiple comparisons. B-D) Quantification of ubiquitination assay performed in HEK293T cells. Briefly, HA-Ubiquitin and FLAG-MECP2 were transfected into cells +/−CDD2021 (1uM) for 48hr followed by an 8hr incubation with (B) DMSO (no inhibitor), (C) proteasome inhibitor (MG-132, 10uM), or (D) lysosome inhibitor (CQ, 40um) before harvest. IP of FLAG in each sample was followed by western blot and quantification of HA (Ubiquitinated MeCP2)/FLAG (total MeCP2). Data presented as mean ± SEM, p-values determined using a Student’s t-test.

**Supplemental Figure 4.**
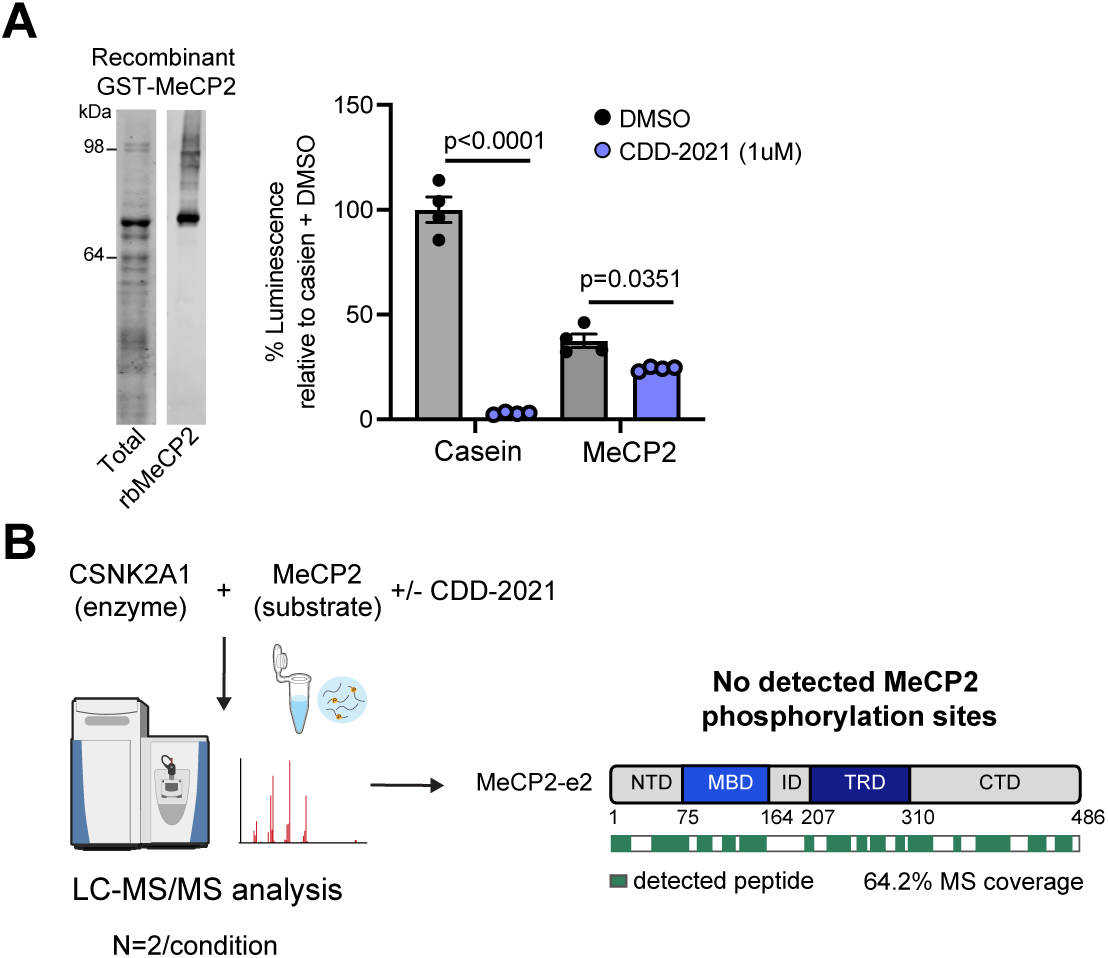
I*n vitro* MeCP2 and CSNK2A1 kinase assay and IP-MS. A) Western blot showing recombinant MeCP2 using total protein stain (left) and MeCP2 antibody (right). ADP-Glo kinase assay using recombinant MeCP2 and CSNK2A1 +/−CDD2021 inhibitor. Phosphorylation activity measured by ATP to ADP conversion to luminescent signal relative to casein + DMSO control group. P-values determined using a two-way ANOVA with post-hoc Sidak’s multiple comparisons test (substrate x treatment interaction, p<0.0001). B) Schematics of IP/MS experiment using *in vitro* recombinant protein assay (10ug of substrate (MeCP2) and 1ug of enzyme (CSNK2A1) with 10uM ATP +/−1uM CDD-2021 or DMSO) and the coverage of human MeCP2-e2.

**Supplemental Figure 5.**
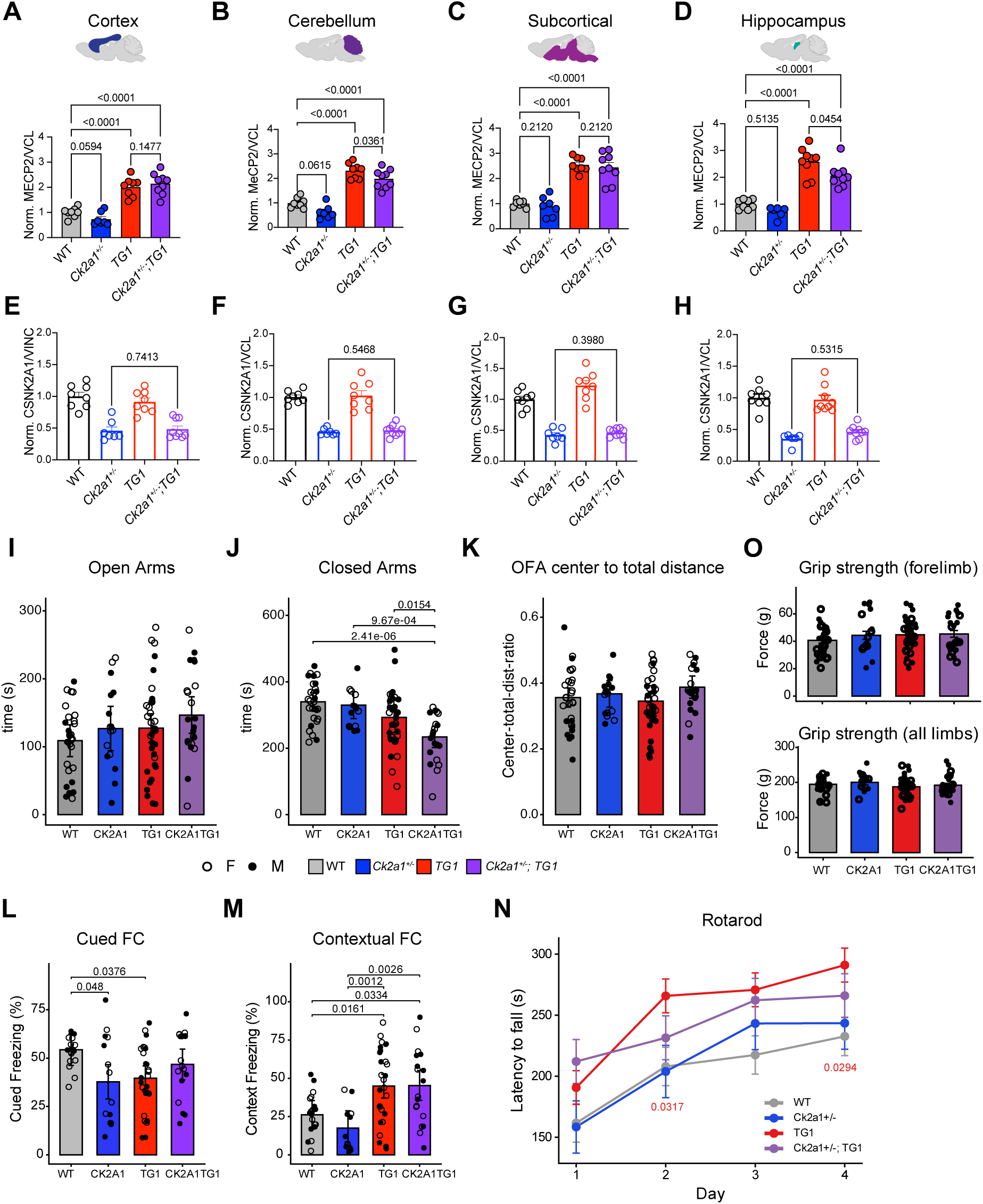
Brain region expression of MeCP2 and CSNK2A1 across *Ck2a1^+/−^* mice crossed to *TG1* mice and behavioral analyses. A-H) Quantification of MeCP2 (A-D) or CSNK2A1 (E-H) levels across all genotypes within distinct brain regions (A,E) cortex, (B,F) cerebellum, (C,G) subcortical, or (D,H) hippocampal samples. P-values determined using a one-way ANOVA with Sidak’s multiple comparisons. I-K) Evaluation of changes in anxiety-related behaviors across all genotypes by examining time spent in (I) open vs (J) closed arms of the elevated plus maze and (K) the center to total distance ratio in the open field assay. L-M) Percentage of time spent freezing in (L) cued and (M) context fear conditioning assay. N) Latency to fall off rotating rod in the rotarod assay and (O) forelimb and all limb grip strength across all genotypes. For EPM, OFA, and rotarod behavior: N=7F, 9M for *Ck2a1^+/−^*, N=10F, 12M for *Ck2a1^+/−^; TG1*, N=16F, 21M for *TG1*, N=19F, 10M for WT. For FC behavior: N=7F, 7M for *Ck2a1^+/−^*, N=9F, 9M for *Ck2a1^+/−^; TG1*, N=13F, 14M for *TG1*, N=13F, 8M for WT. Data presented as mean ± SEM, p-values for behavior tests statistics were determined using a linear model or mixed model (OFA bins) with post-hoc Tukey’s multiple comparisons, see methods for details.

**Supplemental Figure 6.**
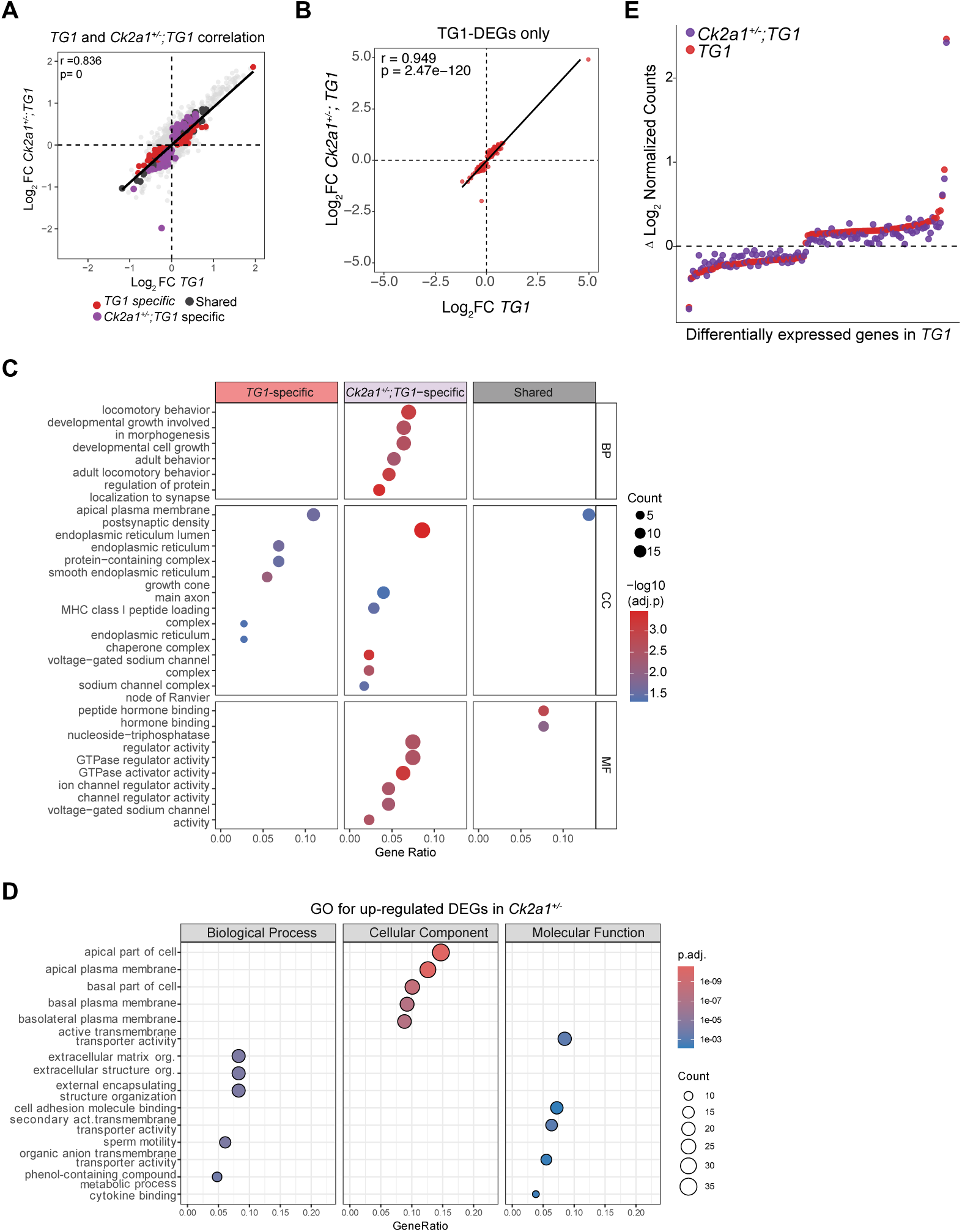
Correlation and GO analyses of transcriptional changes in *Ck2a1^+/−^; TG1* cohort. A) Correlation plot of the Log_2_FC expression values in TG1 and *Ck2a1^+/−^; TG1* samples (red dots are TG1-specific DEGs, purple dots are *Ck2a1^+/−^; TG1* specific DEGs, and dark grey dots are shared DEGs. B) Correlation plot showing high correlation between log_2_FC changes of *TG1*-specific DEGs in both *TG1* and *Ck2a1^+/−^; TG1* samples. C) GO analysis of *TG1*-specific, *Ck2a1^+/−^; TG1*-specific, or shared DEGs. D) GO analysis of *Ck2a1^+/−^* upregulated DEGs. E) Delta Log_2_ normalized counts of *TG1* DEGs in both *TG1* (red) and *Ck2a1^+/−^; TG1* (purple) samples with the dashed line at 0 representing baseline WT levels.

**Supplemental Figure 7.**
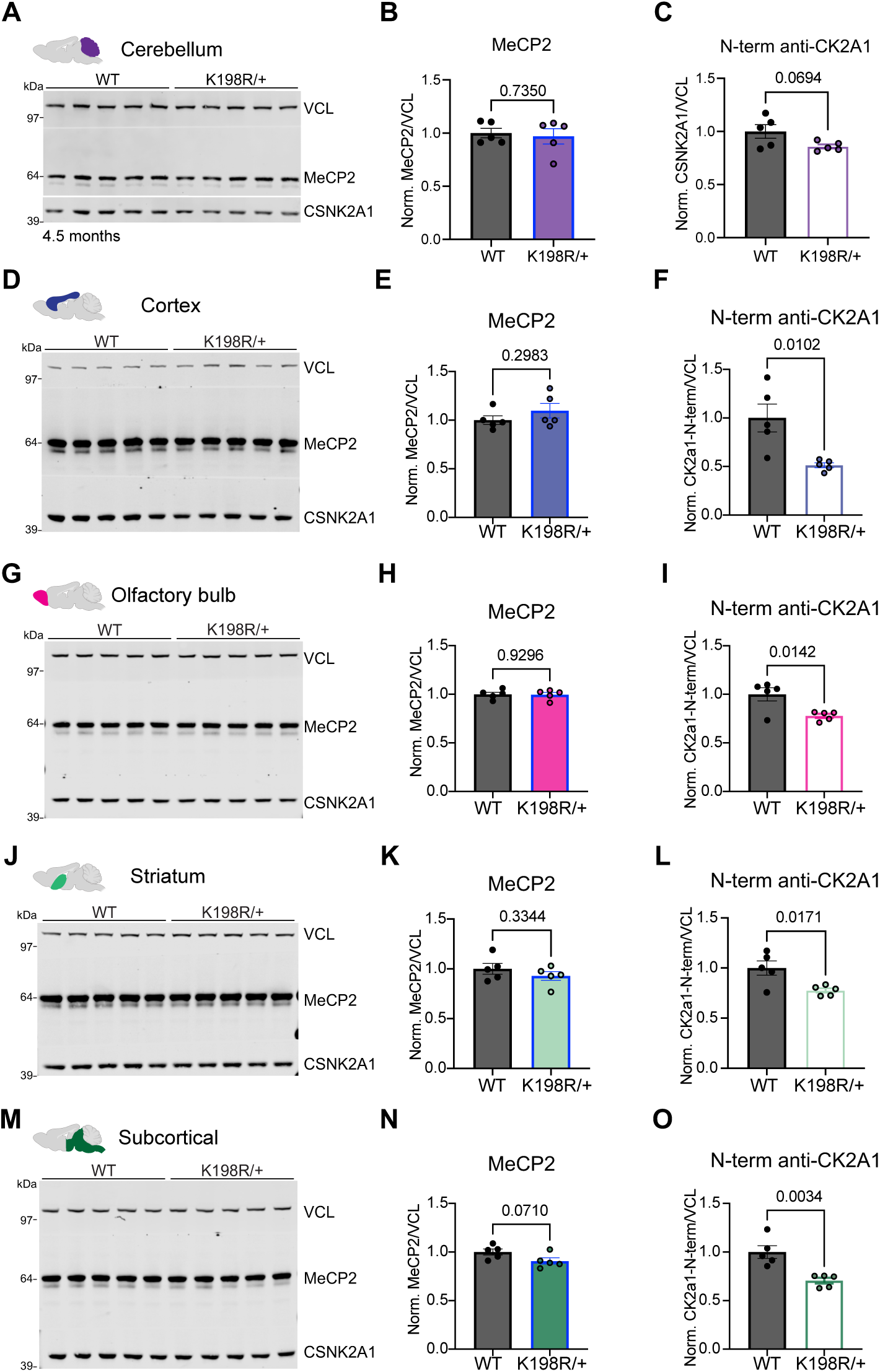
MeCP2 and CSNK2A1 levels in distinct brain regions of *Csnk2a1^K198R/+^*mice. Western blots and quantification for MeCP2 or CSNK2A1 (N-term antibody) relative to loading control (VCL) in distinct brain regions: (A-C) cerebellum, (D-F) cortex, (G-I) olfactory bulb, (J-L) striatum, or (M-O) subcortical tissue. Data presented as mean ± SEM, p-values calculated using a student’s t-test for all WT vs K198R/+ comparisons.

## SUPPLEMENTAL TABLES

**Supplemental Table 1.** CRISPR KO screen results and protein-protein interaction overlap.

**Supplemental Table 2.** Intersection of screen hits with the Drug Gene Interaction Database.

**Supplemental Table 3.** RNA-seq differentially expressed genes results.

**Supplemental Table 4.** Percent rescue of *TG1* and *Ck2a1^+/−^* DEGs in *Ck2a1^+/−^; TG1* samples.

**Supplemental Table 5.** gRNA, shRNA, and primer sequences.

