## Supplemental chemistry methods for "Druggable CSNK2A1-MeCP2 regulatory axis links Okur-Chung neurodevelopmental syndrome with *MECP2*-related disorders"

### Chemistry General Methods:

All starting materials and reagents were purchased from commercial sources and used without further purification. Solvents were purchased as either anhydrous grade products in sealed containers or reagent grade and used as received. All reactions were carried out in dry glassware under a nitrogen atmosphere using standard disposable or gastight syringes, disposable or stainless-steel needles, and septa. Stirring was achieved with magnetic stir bars. Flash column chromatography was performed with SiO2 (230-400 mesh) or by using an automated chromatography instrument with an appropriately sized column. Thin layer chromatography was performed on silica gel 60F254 plates (E. Merck). Non-UV active compounds were visualized on TLC using one of the following stains: KMnO4, ninhydrin, p-anisaldehyde. 1H and 13C NMR spectra were recorded on an instrument operating at either 600MHz, and 151MHz respectively. LCMS data were collected using an HPLC instrument coupled to a low-resolution mass spectrometer with single quadrupole ionization operating in either positive or negative ion mode. The analytical method utilized a C18 column (2.1 × 50 mm, 1.8 μm) eluting with a linear gradient of 95%/5% water/CH3CN (modified with 0.05% formic acid; T = 0 min flow = 0.35 mL/min) to 95%/5% CH3CN/water (T = 3.5 min flow = 0.5 mL/min) then 95%/5% CH3CN/water to T = 5min (0.5 mL/min). Peak detection was done at 254 nm and 230 nm for UV active compounds.

**CSNK2A1 Hit compound**:

DEL selections of CSNK2A1 with Center for Drug Discovery, Baylor College of Medicine's DEL libraries yield the following hit compound from qDOS11 library. [1,2,3]

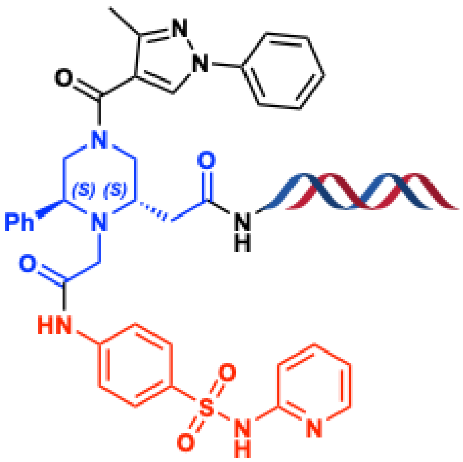


Enrichment profile of qDOS11 hit against CSNK2A1 at 1 μM, observed counts 22970, with the z-score of 436.8. The selections were performed as previously described. [1,2,3]

**Synthesis of CDD-2021:**

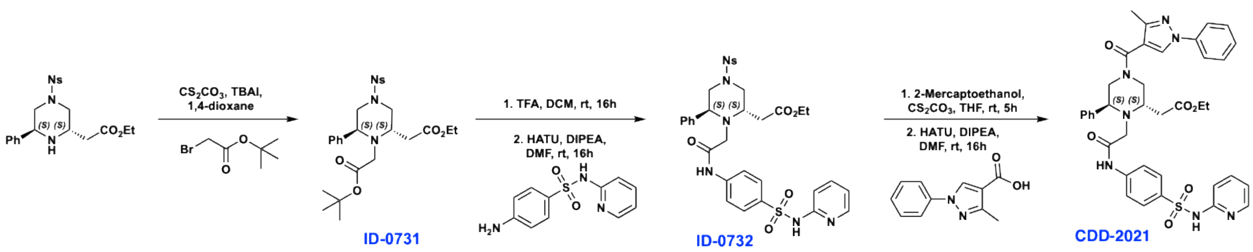

**Experimental procedures and NMR data**.

Note: CDD-2021 was synthesized by following previously reported methods.[1,2,3,4]

**Synthesis of tert-butyl 2-((2S,6S)-2-(2-ethoxy-2-oxoethyl)-4-((4-nitrophenyl)sulfonyl)-6-phenylpiperazin-1-yl)acetate (ID0731).** **$^1$H NMR (600 MHz, $CDCl_3$)** δ 7.83 (dd, *J* = 7.9, 1.6 Hz, 1H), 7.64-7.61 (3, 1H), 7.59-7.57 (m, 1H), 7.54-7.53 (m, 1H), 7.26 – 7.24 (m, 5H), 4.07 (qd, *J* = 7.2, 2.8 Hz, 2H), 3.92 (dd, *J* = 10.8, 3.5 Hz, 1H), 3.78 (dt, *J* = 12.3, 2.2 Hz, 1H), 3.68 (dt, *J* = 12.2, 2.8 Hz, 1H), 3.62 (dd, *J* = 9.3, 3.7 Hz, 1H), 3.20 (d, *J* = 12.2 Hz, 1H), 3.02 (d, *J* = 17.6 Hz, 1H), 2.89 (d, *J* = 17.2 Hz, 1H), 2.81 – 2.70 (m, 2H), 2.65 (dd, *J* = 16.6, 3.6 Hz, 1H), 1.27 (s, 9H), 1.20 (t, *J* = 7.1 Hz, 3H). **$^{13}$C NMR (151 MHz, $CDCl_3$)** δ 172.26, 169.96, 148.44, 133.69, 131.51, 131.22, 131.05, 128.86, 128.51, 128.39, 124.14, 81.02, 77.23, 77.18, 77.02, 60.72, 59.55, 53.66, 52.80, 52.70, 50.24, 29.01, 27.95, 14.17.

**Synthesis of ethyl 2-((2S,6S)-4-((4-nitrophenyl)sulfonyl)-1-(2-oxo-2-((4-(N-(pyridin-2-yl)sulfamoyl)phenyl)amino)ethyl)-6-phenylpiperazin-2-yl)acetate (ID0732).** **$^1$H NMR (600 MHz, $CDCl_3$)** δ 9.34 (s, 1H), 8.32 (d, *J* = 5.8 Hz, 1H), 7.97 (dd, *J* = 7.8, 1.6 Hz, 1H), 7.81 (d, *J* = 8.4 Hz, 2H), 7.75 – 7.65 (m, 6H), 7.44 (d, *J* = 8.8 Hz, 1H), 7.24 (s, 5H), 6.83 (t, *J* = 6.5 Hz, 1H), 4.21 – 4.13 (m, 3H), 3.89 (d, *J* = 12.4 Hz, 1H), 3.77 (d, *J* = 12.8 Hz, 1H), 3.52 (bs, 1H), 3.35 (d, *J* = 12.7 Hz, 1H), 3.20-3.09 (m, 3H), 2.96 – 2.89 (m, 1H), 2.79 (dd, *J* = 16.9, 7.7 Hz, 1H), 1.26 (t, *J* = 7.1 Hz, 3H). **$^{13}$C NMR (151 MHz, $CDCl_3$)** δ 171.81, 154.46, 148.27, 141.81, 134.06, 131.84, 131.28, 129.14, 128.07, 127.87, 124.37, 119.25, 115.03, 61.22, 54.42, 53.19, 14.15.

**Synthesis of ethyl 2-((2S,6S)-4-(3-methyl-1-phenyl-1H-pyrazole-4-carbonyl)-1-(2-oxo-2-((4-(N-(pyridin-2-yl)sulfamoyl)phenyl)amino)ethyl)-6-phenylpiperazin-2-yl)acetate (CDD-2021).** **$^1$H NMR (600 MHz, $CDCl_3$)** δ 9.34 (s, 1H), 8.34 (d, *J* = 5.9 Hz, 1H), 8.01-7.97 (m, 1H), 7.86 (d, *J* = 8.4 Hz, 2H), 7.68-7.65 (m, 3H), 7.62 (d, *J* = 8.0 Hz, 2H), 7.43 (q, *J* = 8.4 Hz, 3H), 7.30 (t, *J* = 7.4 Hz, 1H), 7.32 – 7.16 (m, 5H), 6.81 (t, *J* = 6.5 Hz, 1H), 4.19 – 3.88 (m, 3H), 3.52-3.47 (m, 3H), 3.24 – 3.14 (m, 2H), 2.95 (s, 1H), 2.88 (s, 1H), 2.80-2.78 (m, 1H), 2.67 (bs, 1H), 2.43 (s, 3H), 1.21-1.17 (m, 3H). **$^{13}$C NMR (151 MHz, $CDCl_3$)** δ 171.62, 168.57, 165.27, 162.54, 154.91, 142.05, 141.02, 139.38, 136.66, 129.55, 129.15, 128.91, 128.11, 127.86, 127.41, 127.03, 119.27, 119.23, 115.13, 114.54, 61.27, 54.78, 53.44, 36.48, 31.44, 14.06, 13.02.

**Spectral data for intermediates and CDD-2021:**

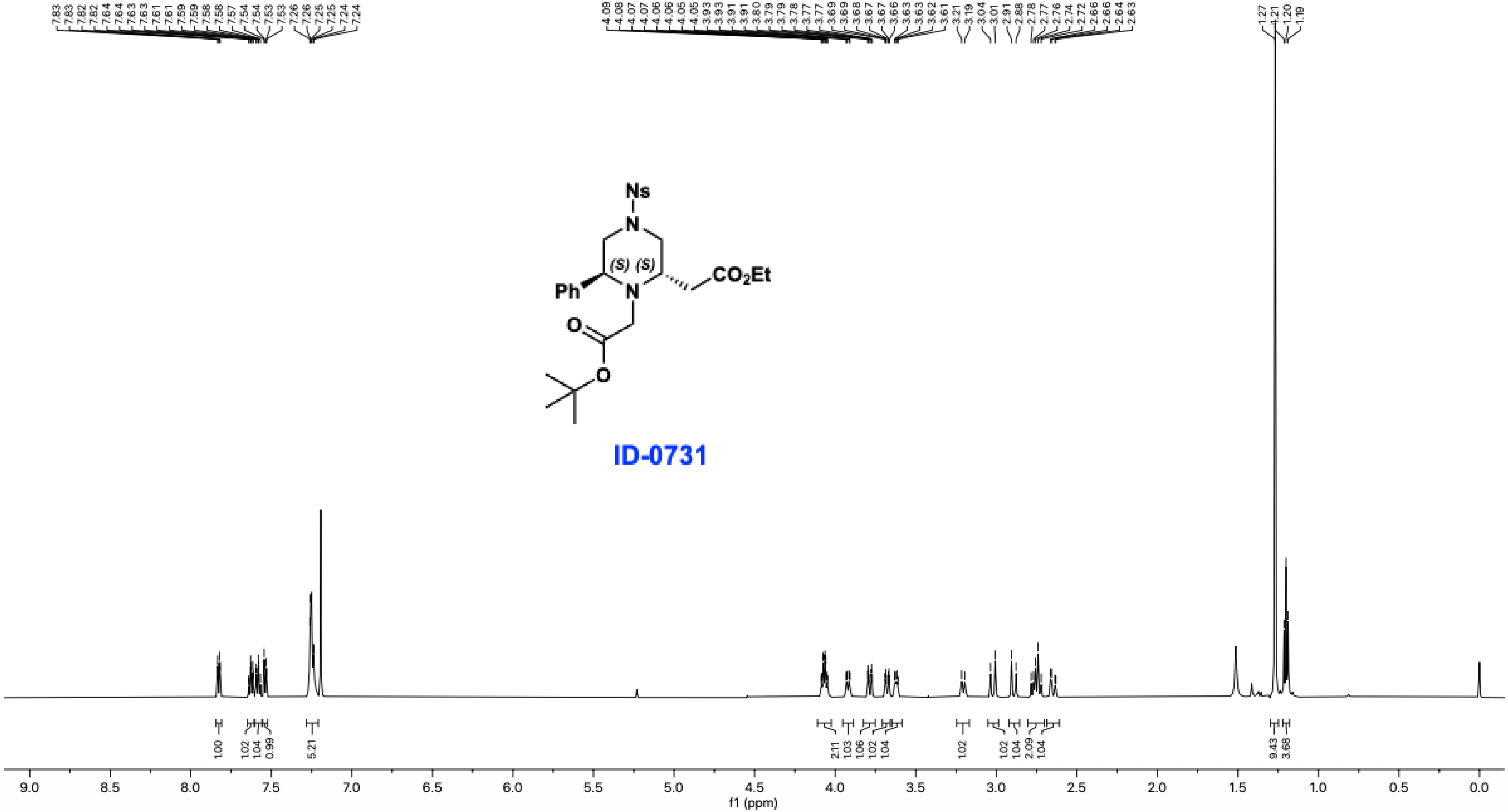

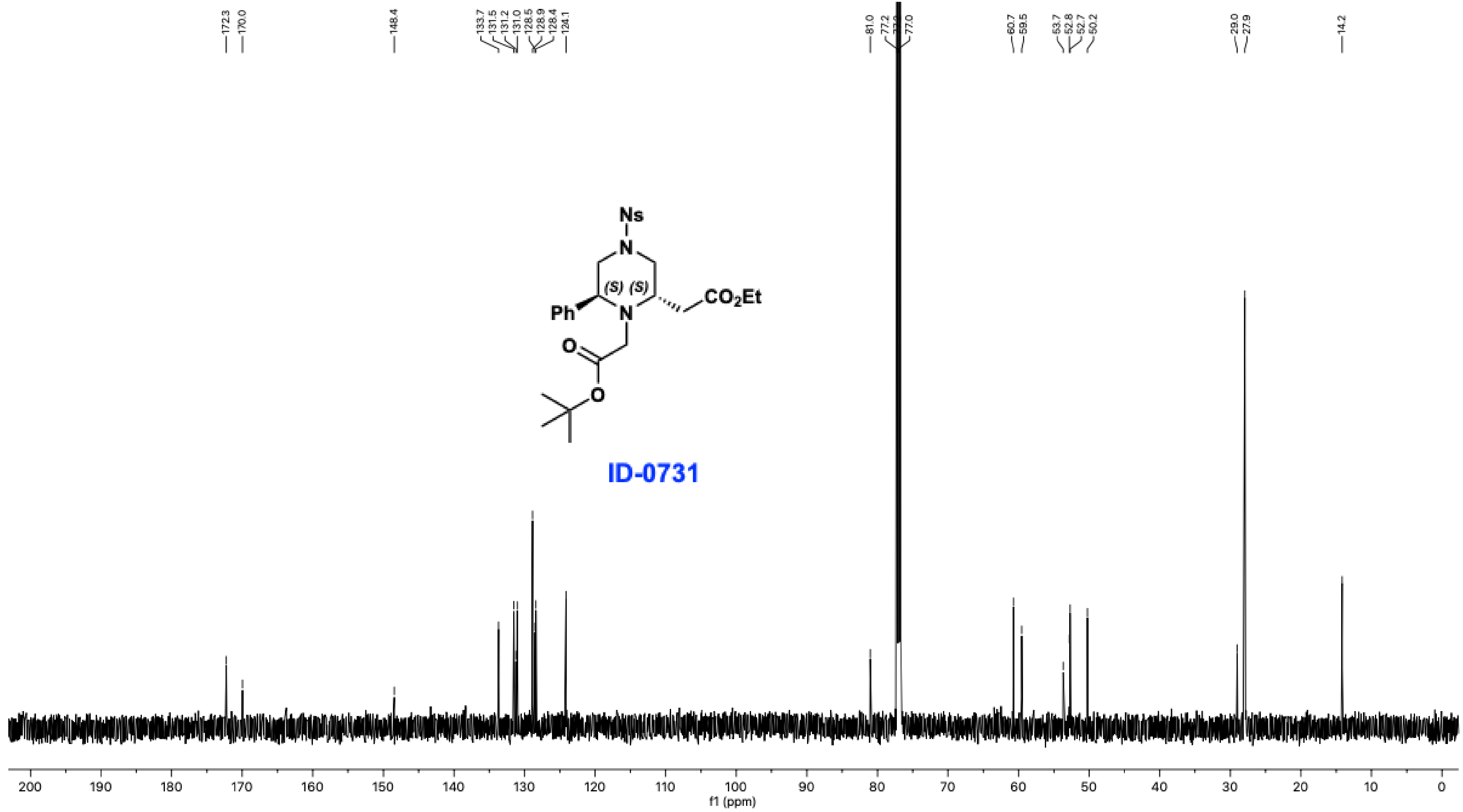

172.3
170.0
148.4
133.7
131.5
131.2
131.0
128.5
128.9
128.4
124.1
81.0
77.2
77.0
60.7
59.5
53.7
52.8
52.7
50.2
29.0
27.9
14.2
Ns
N
(S) (S)
Ph
N
CO2Et
O
O
ID-0731
200 190 180 170 160 150 140 130 120 110 100 90 80 70 60 50 40 30 20 10 0
f1 (ppm)

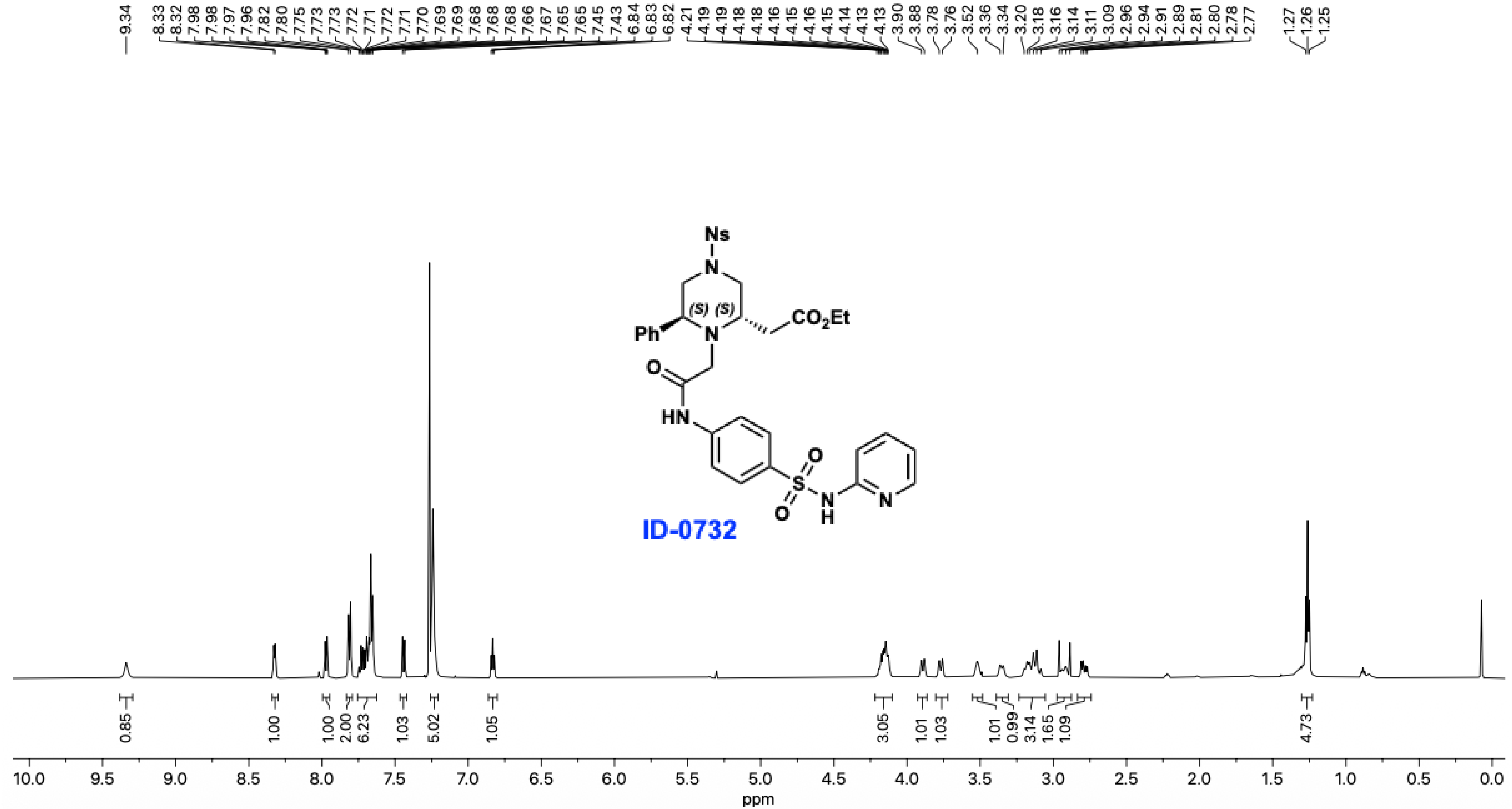

9.34
8.33 8.32 7.98 7.98 7.97 7.96 7.82 7.80 7.75 7.73 7.73 7.72 7.71 7.72 7.71 7.70 7.69 7.69 7.68 7.68 7.68 7.66 7.67 7.65 7.65 7.45 7.43 6.84 6.83 6.82
4.21 4.19 4.19 4.18 4.18 4.16 4.15 4.16 4.15 4.14 4.13 4.13 3.90 3.88 3.78 3.76 3.52 3.36 3.34 3.20 3.18 3.16 3.14 3.11 3.09 2.96 2.94 2.91 2.89 2.81 2.80 2.78 2.77
1.27 1.26 1.25
Ns
N
(S) (S)
Ph
N
CO2Et
O
HN
O
S
O
N
H
N
ID-0732
0.85 1.00 1.00 2.00 6.23 1.03 5.02 1.05 3.05 1.01 1.03 1.01 0.99 3.14 1.65 1.09 4.73
10.0 9.5 9.0 8.5 8.0 7.5 7.0 6.5 6.0 5.5 5.0 4.5 4.0 3.5 3.0 2.5 2.0 1.5 1.0 0.5 0.0
ppm

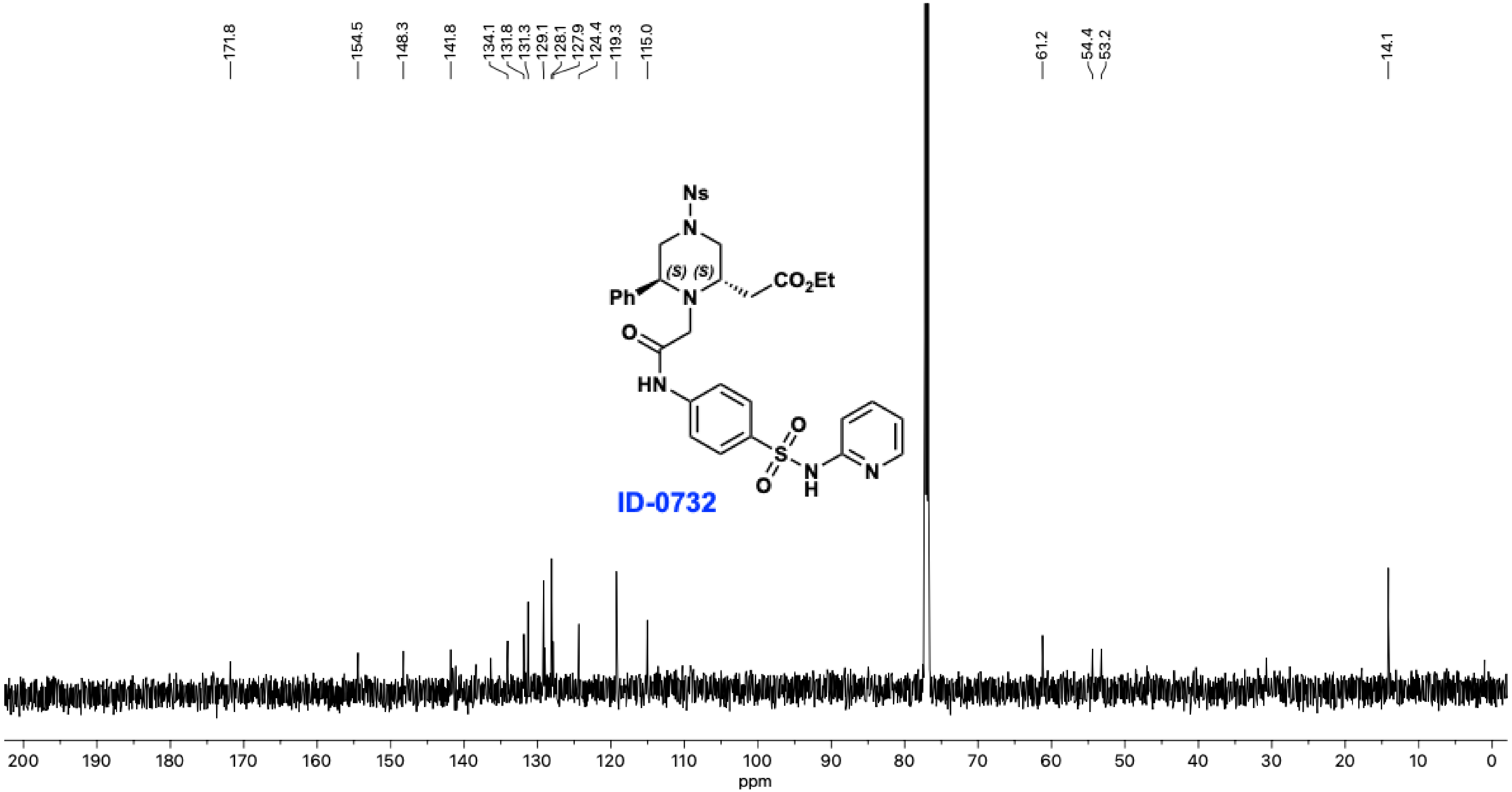
171.8
154.5
148.3
141.8
134.1
131.8
131.3
129.1
128.1
127.9
124.4
119.3
115.0
61.2
54.4
53.2
14.1
Ns
N
(S)
(S)
Ph
N
CO2Et
O
HN
O
S
O
N
H
N
ID-0732
200 190 180 170 160 150 140 130 120 110 100 90 80 70 60 50 40 30 20 10 0
ppm

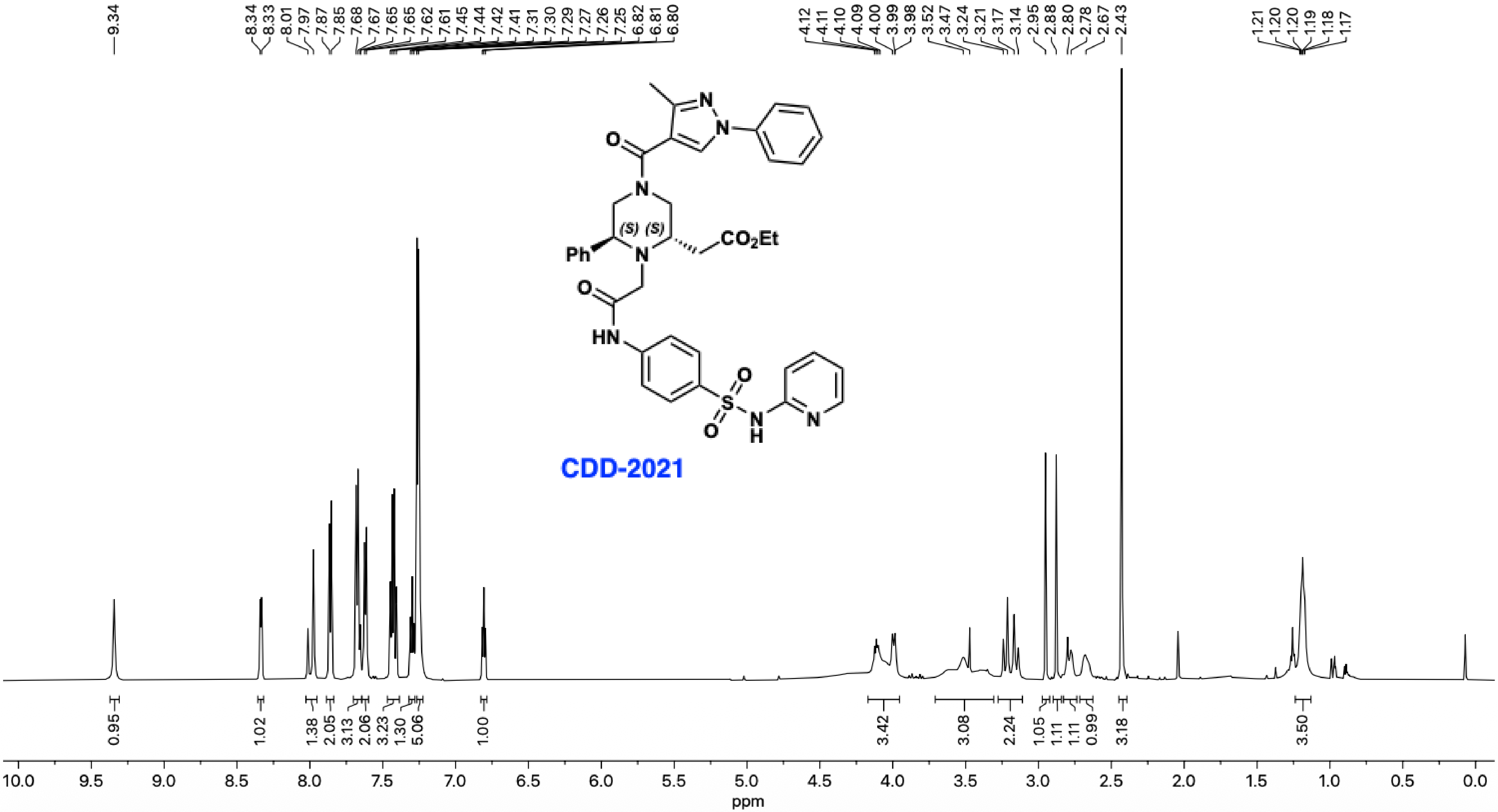
9.34
8.34 8.33 8.01 7.97 7.87 7.85 7.68 7.67 7.65 7.65 7.62 7.61 7.45 7.44 7.42 7.41 7.31 7.30 7.29 7.27 7.26 7.25 6.82 6.81 6.80
4.12 4.11 4.10 4.09 4.00 3.99 3.98 3.52 3.47 3.24 3.21 3.17 3.14 2.95 2.88 2.80 2.78 2.67 2.43
1.21 1.20 1.20 1.19 1.18 1.17
O
N
N
O
N
(S) (S)
Ph
N
CO2Et
O
HN
O
S
O
N
H
N
CDD-2021
0.95 1.02 1.38 2.05 3.13 2.06 3.23 1.30 5.06 1.00 3.42 3.08 2.24 1.05 1.11 1.11 0.99 3.18 3.50
10.0 9.5 9.0 8.5 8.0 7.5 7.0 6.5 6.0 5.5 5.0 4.5 4.0 3.5 3.0 2.5 2.0 1.5 1.0 0.5 0.0
ppm

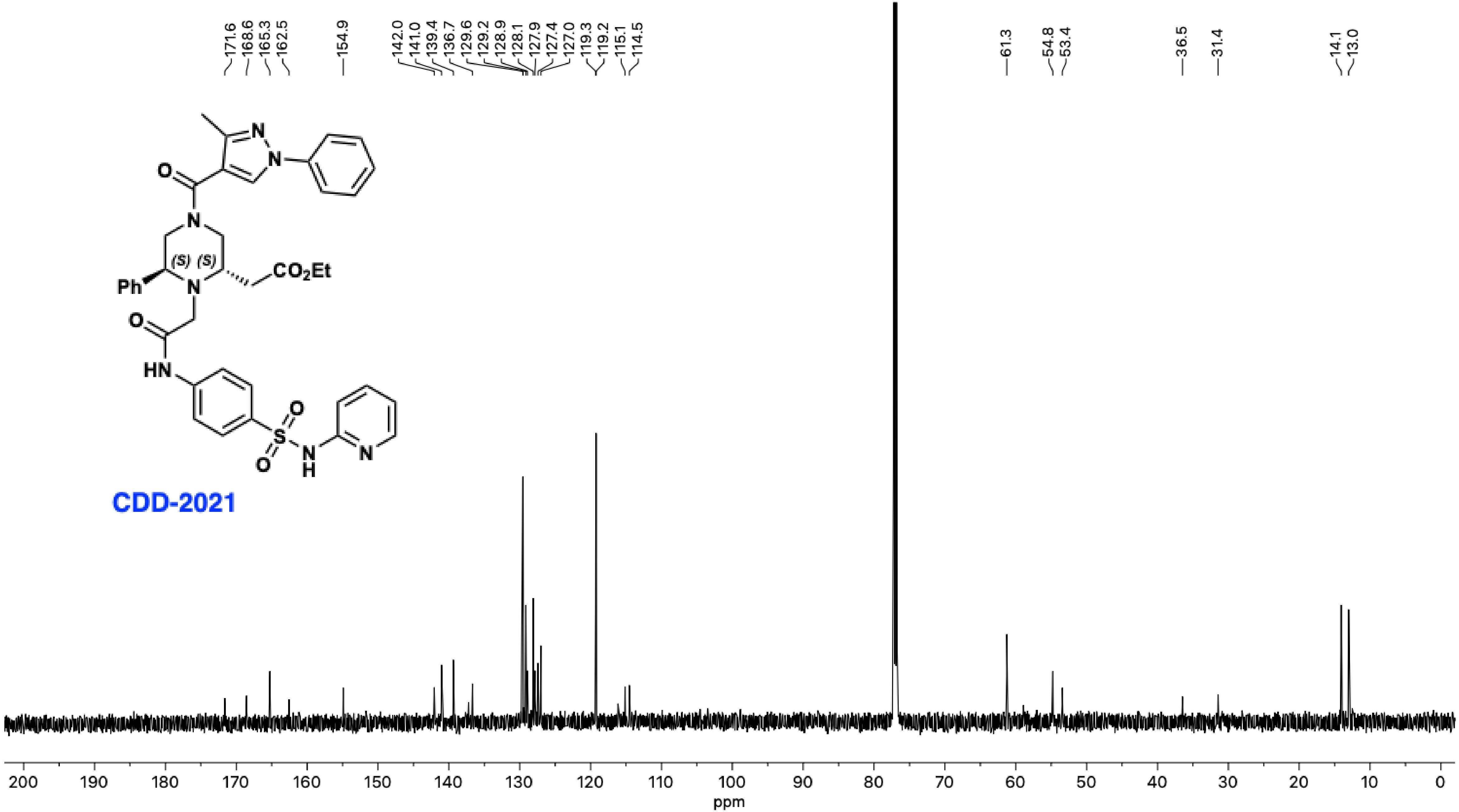
171.6
168.6
165.3
162.5
154.9
142.0
141.0
139.4
136.7
129.6
129.2
128.9
128.1
127.9
127.4
127.0
119.3
119.2
115.1
114.5
61.3
54.8
53.4
36.5
31.4
14.1
13.0
N
N
O
N
(S)
(S)
$CO_2Et$
Ph
N
O
HN
O
S
O
N
H
N
CDD-2021
200
190
180
170
160
150
140
130
120
110
100
90
80
70
60
50
40
30
20
10
0
ppm
